# Transitive interactions among rhizobia determine their symbiotic fitness

**DOI:** 10.64898/2026.01.05.697666

**Authors:** Margarita Granada Agudelo, Bryan Ruiz, Jean-Baptiste Ferdy, Delphine Capela, Philippe Remigi

## Abstract

During host-microbe symbioses, the fitness of mutualistic microbes is determined by the interactions that concurrently occur, throughout their life cycle, with their host and other members of the surrounding microbial community. Disentangling how these multiple interactions shape the fitness of microbial symbionts is challenging, but is essential to understand the diversity and functioning of mutualisms. Here we examined the different fitness components of rhizobial symbionts of the legume plant *Mimosa pudica* across the multiple stages of their symbiotic life cycle. By comparing rhizobial symbiotic fitness in single and pairwise inoculations, we found that inter-bacterial interactions causing significant fitness effects are common, transitive and can have major consequences, sometimes leading to the extinction of a strain. These interactions predominantly occur at the root infection (nodulation) step, but smaller post-infection interaction effects, involving yet uncharacterized mechanisms, were also detected. Furthermore, considering pairwise interactions was sufficient to predict fitness ranks in more complex rhizobial communities consisting of 6 or 8 strains, indicating that higher-order interaction effects do not play a significant role in these communities. Overall, our results provide a quantitative framework to describe the main drivers of rhizobial symbiotic fitness in a simple community context.

## Introduction

Eukaryotic organisms live in tight association with symbiotic microbial communities. In these communities, microbial members, in isolation and in combination with others, can have strong effect on the fitness of their hosts (affecting nutrition, growth, reproduction, health…), along a continuum ranging from deadly pathogens to obligate mutualists [1]. Eukaryotes have thus evolved complex control mechanisms, including multi-layered immune systems, in order to be able to benefit from mutualistic microbes while avoiding pathogens [2]. In return, symbiotic microbes deploy strategies to overcome or endure host control, and to complete their life cycle successfully. The interplay between these infectious strategies and host control mechanisms, which can result from long-term co-evolutionary processes (or from chance in the case of new associations), determines the fitness and evolutionary success of the different organisms involved in these interactions. In practice, however, quantifying the main drivers of microbial fitness is not trivial. Symbiotic microbes, particularly those with a biphasic life cycle (having both an environmental and host-associated phase), are subjected to numerous selective pressures and fitness trade-offs [3–5], and the relative importance of the different microbial traits on fitness is often unclear.

Rhizobia are diazotrophic soil bacteria that are able to engage in mutualistic interactions with legume plants, during which bacteria fix atmospheric nitrogen and provide ammonium to their host plant in exchange for carbon compounds derived from photosynthesis [6]. One hallmark of this symbiosis is the formation of dedicated root organs, called nodules, which house millions of intracellular symbiotic bacteria. These organs are formed following the mutual recognition between the two partners, involving the exchange of chemical signals and subsequent signalling cascades [7]. In most cases, only one bacterium is trapped within the curl of a root hair, which invaginates to form a tubular structure, called infection thread, where bacteria divide and progress towards inner root tissues [8]. At the same time, root tissues at the bases of infected root hair actively divide to form the nodule. Bacteria are then released within nodule cells, where they differentiate into nitrogen-fixing bacteroids. During this symbiosis, mutualism (nitrogen fixation) occurs in the late stages of the interaction and is thus decoupled from the early infection process [9–12]. This creates an opportunity for non-mutualistic bacteria (*e.g*. bacteria that fix little or no nitrogen) to invade legume roots and benefit from the interaction without providing any benefit to their host plant [13].

In rhizobia, symbiotic fitness, which we define here as the number of bacteria detected within plants, depends on three main phenotypic traits: (i) the proliferation and survival of bacteria in the soil and in the rhizosphere, (ii) the ability to induce the formation of new nodules and compete with other rhizobial strains, and (iii) the ability to infect nodule cells, survive within them and fix nitrogen. Rhizobial fitness in a given environment depends not only on the genetic characteristics of the focal strain, but also on the genetic and physiological characteristics of its host plant. Indeed, legume plants employ several mechanisms to monitor and control bacterial infection throughout the symbiotic interaction [14–16]. These mechanisms can be classified depending on whether they act during pre-or post-infection processes. Pre-infection control mechanisms mainly refer to the initiation of nodulation by rhizobia from the rhizosphere. These mechanisms rely on mutual recognition processes between partners and involves on the bacterial side the production of Nod factors, surface polysaccharides or secreted effectors [17]. The ability of a focal strain to form nodules on its host plant when competing with other strains present in the rhizosphere (*i.e.* nodulation competitiveness) involves a wider range of bacterial traits, including motility, stress resistance and metabolic properties. This phenomenon is well-known for explaining the dominance of certain strains in legume nodules [18,19]. At the post-infection level, bacterial proliferation and survival can be modulated by sanctions and/or rewards, in the form of restricted or preferential nutrient allocation imposed by plants on bacteria, depending on the amount of fixed nitrogen provided by the different strains [14,19–21]. Recently, a mechanism of ‘conditional sanctions’ was described in pea, during which plants allocate more resources to an intermediate fixer when it is co-inoculated with a non-fixer, than when the same strain is co-inoculated with a good fixer [22]. In addition, plant immunity can act within nodules to influence bacterial proliferation and survival [23].

Beyond direct plant-rhizobia interactions, bacterial fitness is also influenced by interactions with other members of the microbiome [24,25]. These interactions can be direct (microbes competing or interfering with each other in the rhizosphere or within nodules) or indirect (e.g., mediated by the host plant during the nodulation or post-infection steps), and may occur at any of the different steps of the rhizobial lifecycle. While the existence of direct and indirect pre-infection competitive effects affecting nodulation have long been recognized [25,26], the analysis of post-infection inter-bacterial interactions has been more sporadic [22], probably because of difficulties to quantify bacterial abundance within individual nodules [25]. Yet, culture-independent methods based on quantitative PCR or deep DNA sequencing [27,28] offer tools for precise quantification of the relative influence of the different components of rhizobial fitness during symbiosis and with different levels of microbial community complexities.

Here we quantified the effect of inter-bacterial interactions on rhizobial abundance during the pre-and post-infection stages of symbiosis. We used the tropical legume plant *Mimosa pudica* and a collection of 9 of its rhizobial symbionts (spanning 2 genera and 4 species of beta-rhizobia) as model species. We first compared rhizobial abundances in single and pairwise inoculation experiments to identify cases where inter-bacterial interactions affect fitness. We then investigated at which stage of the lifecycle these interactions take place. Finally, we tested whether interactions are conserved in more complex communities of 6 and 8 strains, constituting intra-and inter-specific strain assemblages, respectively. Our results showed that nodulation competitiveness is the major type of inter-bacterial interactions driving rhizobial fitness, with post-infection interactions making only minor contributions. Moreover, we observed that the competitive effects in pairwise inoculations were transitive, enabling the prediction of dominance ranks in more complex communities.

## Results

### - Symbiotic fitness of nine rhizobial symbionts of *Mimosa pudica* in single inoculations

To investigate the effect of bacterial interactions on rhizobial fitness, we used a collection of 9 strains that were isolated from different host plants and countries (Table 1, Supplementary Fig. 1). This collection includes 6 *Cupriavidus taiwanensis* strains (abbreviated Ctai1-Ctai6 in the rest of the manuscript) to cover the whole genetic diversity of the *C. taiwanensis* species complex [29]. In addition, we used *Cupriavidus sp.* amp6 (Csp), and 2 *Paraburkholderia* strains: *P. phymatum* STM815 (Pphy) and *P. caribensis* Tj182 (Pcar). All strains have a similar growth rate in rich medium, with a slight reduction observed for the 2 *Paraburkholderia* strains (Supplementary Fig. 2).

**Table 1:** List of strains used in this study.

| STRAIN ID | DESCRIPTION | ORIGIN | HOST PLANT | GENOME SIZE (BP) | REFERENCE |
| --- | --- | --- | --- | --- | --- |
| **Ctai1** | *Cupriavidus taiwanensis* LMG19424, Sm<sup>R</sup> | Taiwan | *Mimosa pudica* | 6,476,523 | [79] |
| **Ctai2** | *Cupriavidus taiwanensis* STM6021 (M1-3C2) | French Guiana | *Mimosa pudica* | 6,811,355 | [43] |
| **Ctai3** | *Cupriavidus taiwanensis* STM8959 (ip2.30) | Philippines | *Mimosa diplotrichia* | 6,705,804 | [80] |
| **Ctai4** | *Cupriavidus taiwanensis* STM8969 (MAPUD 10.1) | Costa Rica | *Mimosa pudica* | 6,760,360 | [81] |
| **Ctai5** | *Cupriavidus taiwanensis* STM8970 (cmp52) | Texas | *Mimosa asperata* | 7,026,539 | [82] |
| **Ctai6** | *Cupriavidus taiwanensis* STM6160 | New Caledonia | *Mimosa pudica* | 7,023,887 | [45] |
| **Csp** | *Cupriavidus sp.* STM8968 (AMP6) | Texas | *Mimosa asperata* | 7,579,563 | [82] |
| **Pphy** | *Paraurkholderia phymatum* STM815 | French Guiana | *Machaerium lunatum* | 8,598,258 | [83] |
| **Pcar** | *Paraburkholderia caribensis* TJ182 | Taiwan | *Mimosa diplotrichia* | 9,205,673 | [68] |
| **Ctai1 $\Delta nifH$** | *nifH* deletion mutant from Ctai1 |  |  |  | [11] |

We first assayed the symbiotic phenotypes of each strain in single inoculation experiments. We inoculated the 9 strains individually on *M. pudica* seedlings and measured the dry weight (DW) of aerial parts, as a proxy for plant fitness, and the number of nodules formed on the entire root system at 28 days post-inoculation. The tested strains formed, on average, between 8.5 (for Ctai2) and 30 (for Pcar) nodules per plant at 28 dpi (Fig. 1A). Almost all strains provided significant and comparable growth benefits to *M. pudica*, with average dry weights ranging from 30 (Ctai5) to 33.6 mg (Ctai4) per plant (Fig. 1B). The only exception is the strain *Cupriavidus sp.* amp6 (Csp) for which plants’ DW was only slightly higher than non-inoculated plants (17.6 vs. 13.9 mg per plant for non-inoculated plants). The symbiotic fitness of bacteria was estimated by measuring the total bacterial load contained within all nodules formed on one plant by quantitative PCR (qPCR) using strain-specific primers (Supplementary Fig. 3; Supplementary Table 1). The median bacterial load values were continuously distributed between approximatively 7×10^7^ (Pcar and Csp) and 2.5×10^8^ (Ctai1) copies of bacterial genomes per plant (Fig. 1C). Overall, these results show that all strains effectively nodulate *M. pudica* and all, with the exception of Csp, are efficient mutualistic symbionts of this plant.

**Figure 1.**
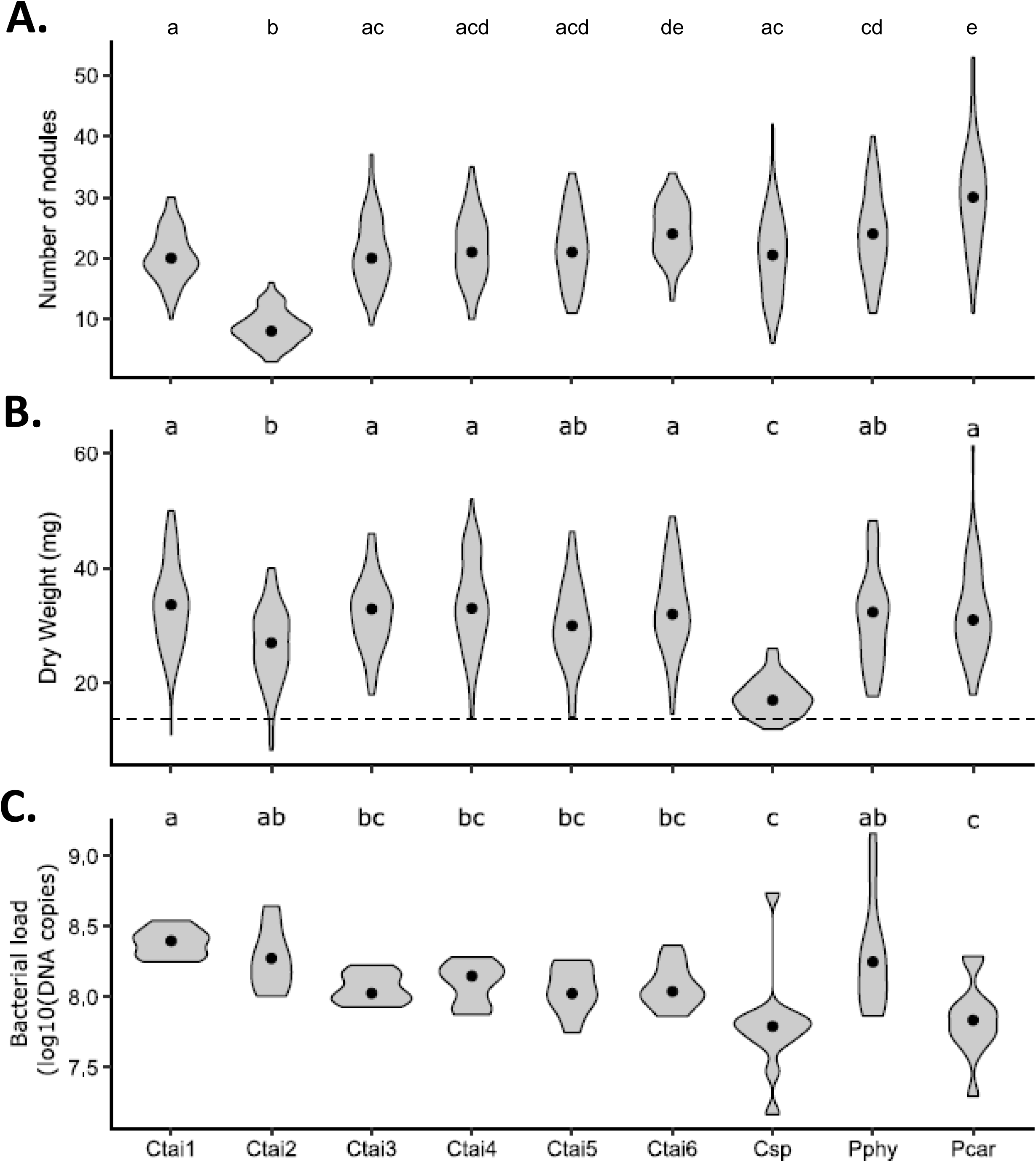
Symbiotic phenotypes of *Mimosa pudica* symbionts in single inoculation. (A) Number of nodules. (B) Plant dry weight. (C) Bacterial load per plant. Measurements were performed on plants inoculated with the different bacterial strains at 28 dpi across four independent experiments. The distributions of data points are shown in grey, black points show the medians. n > 60 (A, B) or n = 12 (C). In (B), the dotted line indicates average dry weight for non-inoculated plants. Statistical analyses were performed with non-parametric Kruskall-Wallis with Dunn’s post-hoc test. Letter groups indicate statistical significance (p < 0.05).

### - Pairwise inoculation experiments reveal a high prevalence of inter-bacterial negative interactions

To detect possible inter-bacterial interactions occurring during symbiosis within our strain collection, we performed 36 pairwise co-inoculations, representing all possible combinations of the 9 strains, and measured the same symbiotic properties as before (number of nodules, plant dry weight and bacterial abundance within nodules). Overall, no systematic bias on plant growth could be detected in pairwise compared to single inoculation experiments (Supplementary Fig. 4), showing that bacterial competition has negligible effects on plant development. However, comparing bacterial abundances in these two settings revealed numerous cases where bacterial loads were different from those expected from single inoculation data (Fig. 2A and Material and methods). We considered the bacterial load to be significantly different in pairwise compared to single inoculation experiments when all the replicate data points from the pairwise comparisons fell outside of the 95% confidence interval predicted from the single inoculation data. In other words, bacterial loads were not deemed as different when at least one pairwise data point fell within the 95% confidence interval. This strict definition was adopted to increase the robustness (while decreasing sensitivity) of our screen aiming at identifying robust, large-effect inter-bacterial interactions affecting fitness. For each pair of strains, we then classified the outcome of inoculation in 3 categories: (i) co-existence of the two strains without interaction, when both strains’ load was not different from single inoculation, (ii) co-existence with interaction, when the abundance of at least one strain was (positively or negatively) affected by the presence of the other strain, and (iii) interaction with exclusion, when the abundance of one strain fell below the detection limit in all replicates. Using this classification, evidence for inter-strain interactions was found in 92% of the cases (33 out of 36 pairs of strains; Fig. 2B). Out of these 33 cases, extinction of one strain was observed in 15 pairs. These results show that interactions between rhizobial strains are frequent and can have major effects, by leading to the extinction of one strain. In the 18 other cases of interactions, both strains remained in co-existence.

**Figure 2.**
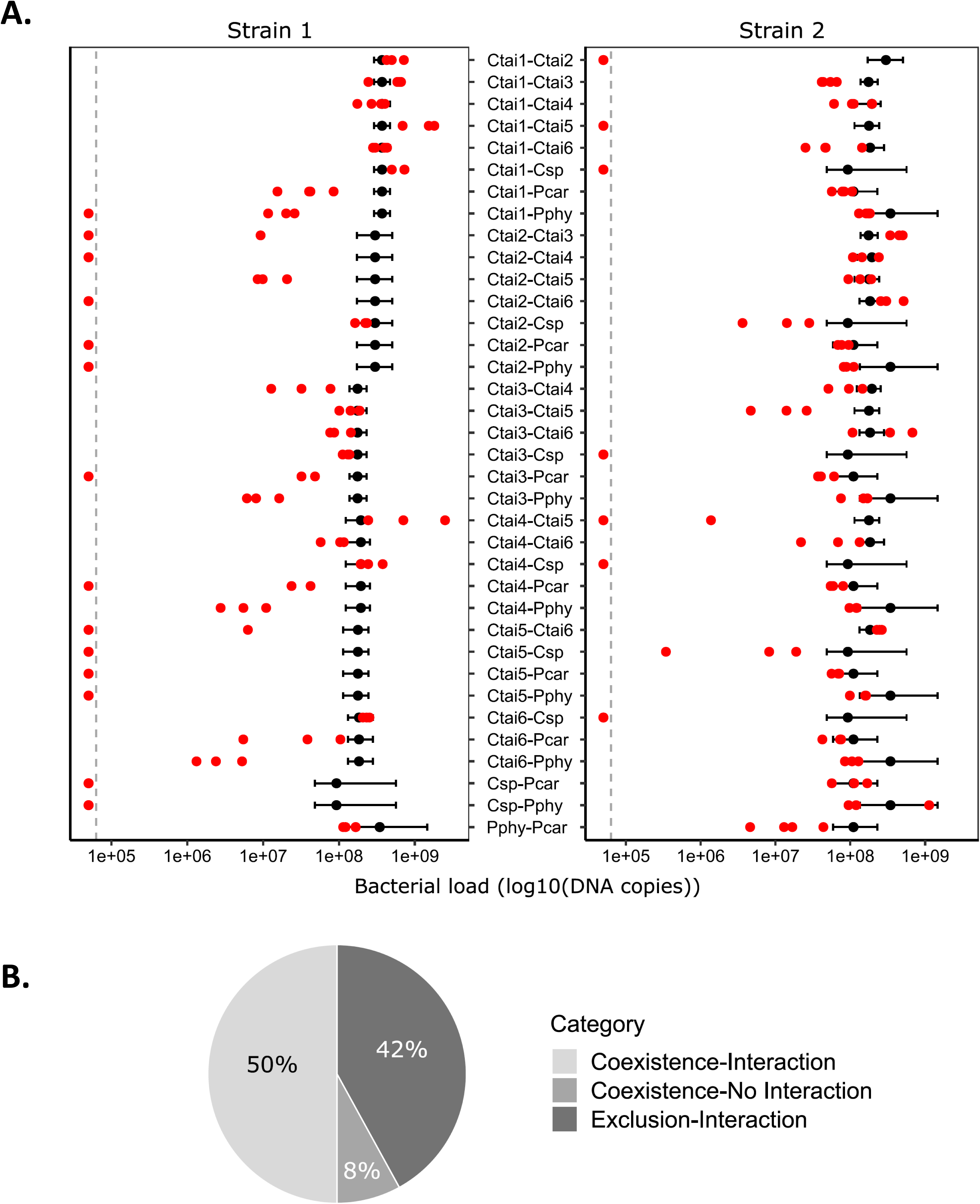
Identification of interactions during symbiosis for 36 pairs of rhizobial strains. (A) Comparison of bacterial loads in nodules from single and pairwise inoculations. Red dots show pairwise inoculation data (n = 3 or 4). Black dots and error bars represent the mean and 95% confidence intervals, respectively, of bacterial loads predicted from single inoculations. The dotted line indicates the limit of detection. (B) Summary of the classification of pairwise inoculations according to the existence and type of inter-bacterial interaction.

### - Inter-strain interactions predominantly occur at the nodulation step

We then sought to identify the step of the symbiotic life cycle at which these interactions occur. In our simplified experimental system, symbiotic fitness of a focal rhizobial strain is dependent on both the number of nodules formed by this strain on a plant and the number of bacteria per nodule. In the pairwise inoculation tests described above, the 15 cases of interactions where one strain was completely excluded were likely due to the strong nodulation competitiveness of the remaining strain, as we calculated that even one nodule colonised by one strain could have been detected by qPCR (Material and Methods).

To gather a more complete view on the role of nodulation competitiveness in our experiments, we also assayed nodulation competitiveness on a selection of 10 pairs of strains showing co-existence in pairwise inoculations (Fig. 3A). Among these 10 pairs, 8 had been classified as cases of interactions, while the 2 others (Ctai1-Ctai4 and Ctai1-Ctai6) showed similar fitness and were used as controls. In one of these control pairs, Ctai1 had slightly higher nodulation competitiveness than Ctai4, challenging the classification of Ctai1-Ctai4 as a ‘non-interacting’ pair. Among the 8 pairs of interacting strains, statistically-significant differences in nodulation competitiveness were observed 6 times. In all these cases, the strain with the highest nodulation competitiveness also had highest relative fitness (Fig. 2A, 3A), suggesting that nodulation competitiveness contributes to the observed unequal fitness. In the remaining 2 cases (Ctai4-Ctai6 and Pphy-Pcar), both strains had similar nodulation competitiveness while they were found to have different fitness. We note that, in both pairs, strains showed only slightly fitness differences in single vs. pairwise inoculations (Fig. 2A), and nodulation competitiveness values were highly variable (Fig. 3A), preventing any strong conclusion to be drawn regarding interactions occurring within these two pairs of strains. Overall differences in nodulation competitiveness likely explain at least 63% of rhizobial interactions detected in pairwise inoculations (21 cases out of 33).

**Figure 3.**
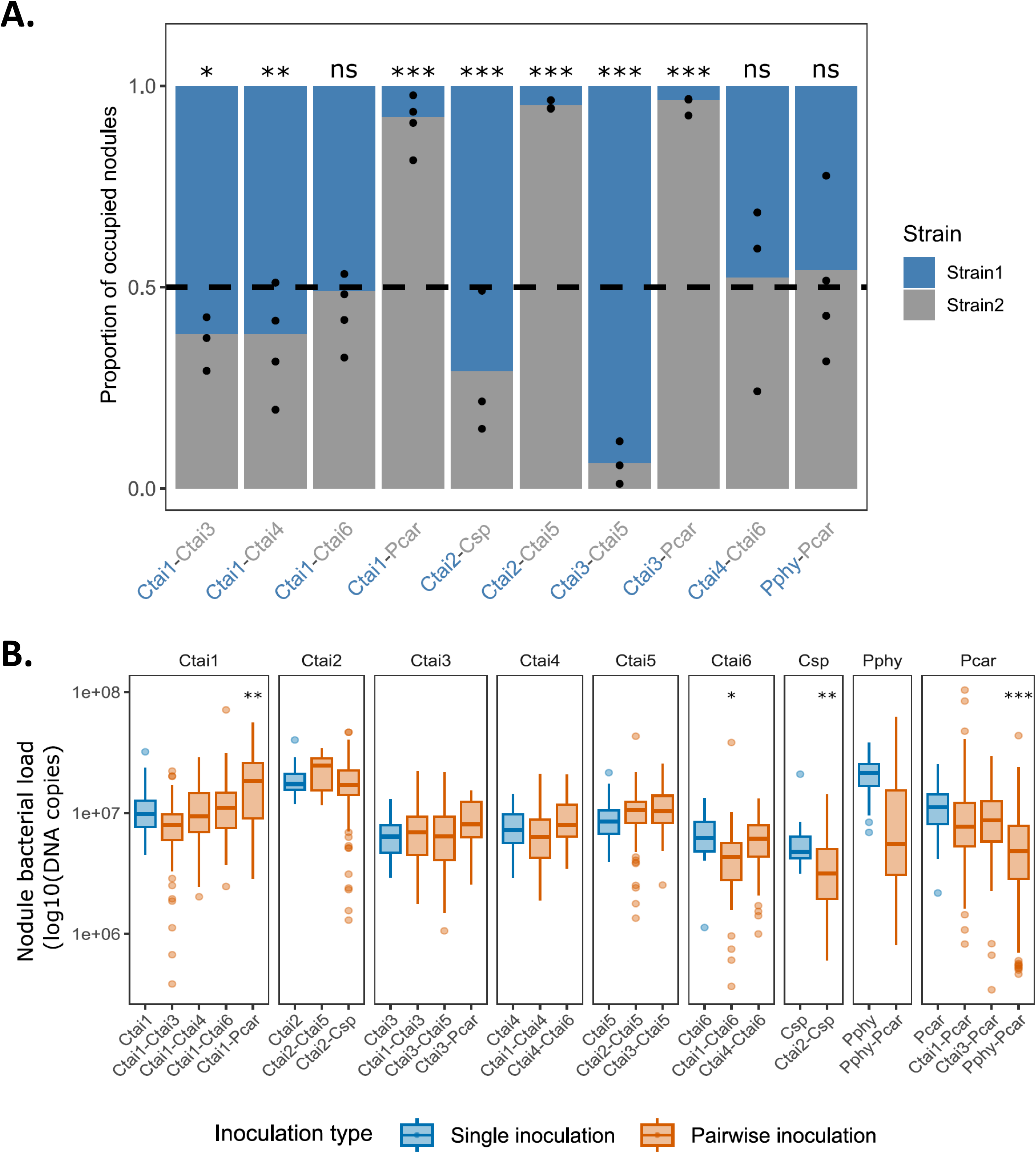
Characterisation of interactions across different steps of the symbiotic life cycle for 10 pairs of strains. (A) Proportion of nodules occupied by each competing strain in pairwise inoculation. Asterisks indicate significant deviations from the expected 50% (equal occupancy) in co-inoculation experiments, reflecting differences in nodulation competitiveness between strains. Statistical significance was assessed using a GLMM with a binomial distribution and logit link function. Detailed statistical results are provided in section 1 of the Supplementary Online Material (SOM). (B) Comparison of bacterial load within individual nodules from single and co-inoculated plants. Each panel shows data for the quantification of one focal strain (indicated above each panel) and represents data from plants inoculated only with this focal strain (blue boxplots) or co-inoculated with another strain (orange boxplots, with the identity of co-inoculated strains indicated below the plots). Statistical significance was assessed using a GLM with a Gamma distribution and logarithmic link function. Detailed statistical results are provided in section 2 of SOM. On both plots, *, ** and *** indicate a p-value below 0.05, 0.01 or 0.001, respectively.

Variations in nodulation competitiveness can be caused either by differential growth or survival in the rhizosphere, and/or by preferential compatibility between symbiotic partners at the early stages of the interaction. To investigate which of these two mechanisms might be responsible for the effects observed above, rhizospheric bacterial populations were measured in single and pairwise inoculations at 7 days post-inoculation, when the first nodules became visible. In single inoculations, median bacterial loads in the rhizosphere varied between 10^5^ to 5×10^6^ DNA copies per ml of medium (Supplementary Fig. 5A). Except for Pcar, strains with high nodulation competitiveness (*e.g*. Pphy, Ctai1) did not have a markedly improved survival in the rhizosphere and, vice versa, some poor competitors (*e.g.* Ctai2) had a relatively high survival in the rhizosphere. However, in pairwise inoculations, a significant positive correlation was found between the proportion of bacterial loads in the rhizosphere and the proportion of occupied nodules (Supplementary Fig. 5B; Spearman correlation test, rho =0.67, p = 4.4×10^-6^). The overall correlation between these two traits suggests that high competitiveness in the rhizosphere might underpin high nodulation competitiveness for a number of bacterial pairs. However, significant outliers were also detected, with strains showing high nodulation competitiveness despite being outcompeted in the rhizosphere. In these cases, the effect of survival in the rhizosphere is likely insignificant, and the difference in nodulation competitiveness between the two strains must instead be linked to direct plant-bacteria signalling compatibility.

### - Evidence for post-infection control mechanisms

So far, we have analysed pre-infection competition effects occurring before the establishment of an effective mutualistic symbiosis. Once mature nodules are formed, post-infection interactions may also contribute to modify rhizobial fitness during pairwise inoculations. The pairwise nodulation competitiveness assays described previously also allowed us to measure bacterial abundances in individual nodules, and compare these values to those found in single inoculations (Fig. 3B). In most cases, bacterial abundances in individual nodules were similar in nodules arising from single-or pairwise-inoculated cases. However, significant differences were observed in four cases. The abundance of Ctai1 increased when co-inoculated with Pcar. In the remaining three cases, co-inoculation led to a decrease in the abundance of one strain (Ctai6, Csp and Pcar, when co-inoculated with Ctai1, Ctai2 and Pphy, respectively). Interestingly, Csp is a poor mutualist compared to Ctai2 (Fig. 1B). Therefore, the decrease in Csp abundance during co-inoculation with Ctai2 is consistent with the mechanism of conditional sanctions previously described in pea, where the host plant can modulate bacterial fitness within nodules during pairwise inoculations depending on the relative benefit that each strain provides to its host [22].

To further investigate conditional post-infection control in *M. pudica*, we focused on the Ctai1-Ctai6 pair since the fitness and nodulation measurements for these strains showed low variability, and their equal nodulation competitiveness facilitates the recovery of individual nodules from each strain when performing fitness measures in pairwise inoculations. In this pair of strains, Ctai1 caused a moderate (less than two-fold) reduction in Ctai6 load (Fig. 3B). The interaction between Ctai1 and Ctai6 seems specific, since Ctai1 did not cause a reduction in the load of Ctai3 or Ctai4, and the load of Ctai6 was not reduced when co-inoculated with Ctai4. Another set of experiments was performed to confirm and further explore this phenomenon. To test if conditional sanctions could also operate in this case, Ctai6 was co-inoculated with a non-fixing *ΔnifH* mutant of Ctai1 [11]. A similar decrease in Ctai6 abundance was observed when it was co-inoculated with the *nifH* mutant or with the WT Ctai1 strains (Fig. 4A), showing that this post-infection interaction effect was not linked to the nitrogen fixation level of the competing strain. Moreover, to confirm that the reduction in Ctai6 bacterial load, as measured by qPCR, indeed corresponds to a reduction in the number of viable bacteria, we measured the number of colony forming units (CFU) in individual nodules from single or co-inoculated plants (Fig. 4B). We observed that the median number of viable Ctai6 bacteria in nodules was approximately 10 times lower during pairwise inoculation with Ctai1 than when inoculated alone or co-inoculated with Ctai4, showing that qPCR data, probably by detecting DNA of non-viable bacteria, underestimate the viability loss of rhizobia within nodules. These results confirm the specific and marked reduction in Ctai6 fitness during pairwise inoculation with Ctai1.

**Figure 4.**
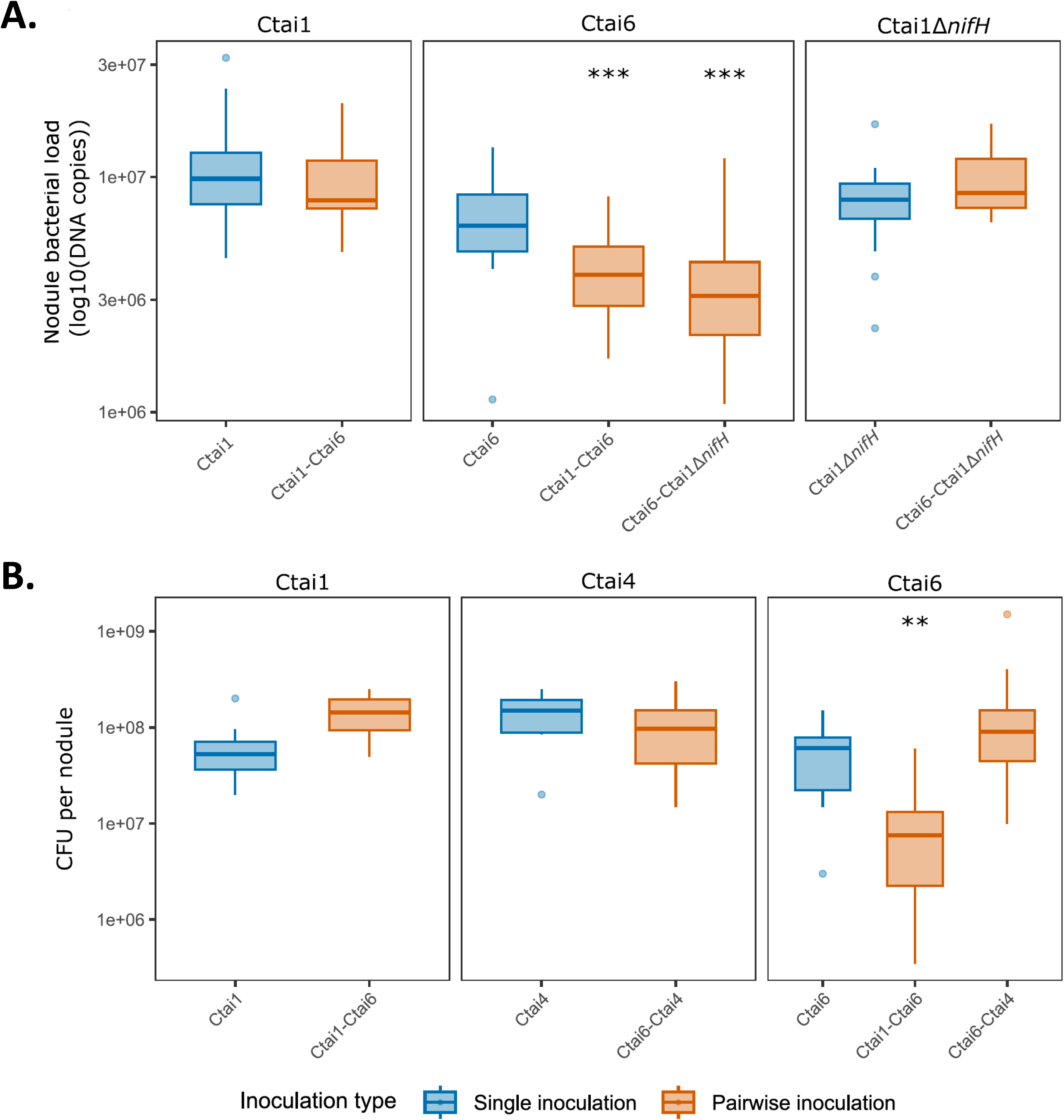
A specific post-infection interaction between Ctai1 and Ctai6 decreases fitness of Ctai6 independently of nitrogen fixation. (A) Bacterial fitness in individual nodules evaluated by qPCR. (B) Bacterial fitness in individual nodules evaluated by dilution plating (CFU counts). Each panel shows data for the quantification of one focal strain (indicated above each panel) and represents data from plants inoculated only with this focal strain (blue boxplots) or co-inoculated with another strain (orange boxplots, identity of co-inoculated strains indicated below the plots). Statistical significance was assessed using a Student’s t-test. On both plots, ** and *** indicate a p-value below 0.01 or 0.001, respectively. In (A), n ≥ 27 except for Ctai1 Δ*nifH* in the Ctai6-Ctai Δ*nifH* mix where n = 3. In (B), n= 10 for single inoculations and n ≥ 4 for pairwise inoculations.

### - Transitivity in rhizobial interactions is conserved in more complex communities

A current challenge in microbial ecology is to assess to which extent the results of pairwise interaction studies are relevant for communities comprising numerous strains or species. Indeed, pairwise interactions often fail to predict strain abundance at a given time point in more complex microbial communities due to the existence of intransitive or higher-order interactions [30,31]. To test for the existence of intransitive interactions, we first re-analysed bacterial fitness data from pairwise inoculation tests (shown in Fig. 2) by analysing, across all experiments, the frequency at which each strain was more abundant than its competitor (Fig. 5A). We found that the network of interactions only consists of transitive interactions, even if competition outcomes were occasionally inverted in some individual replicates of given pairs (e.g. Ctai3-Pcar, Ctai6-Pcar, Ctai1-Ctai4, Ctai3-Ctai6 and Ctai4-Ctai6). This result prompted us to test if this dominance hierarchy was conserved in communities comprising more than 2 members. To do so, we first computed a ‘dominance index’, ranking strains according to their average dominance in pairwise competitions (Fig. 5B, see Material and Methods for details on index calculation). Next, we experimentally measured relative strains’ fitness in a community comprising 8 strains (Fig. 5C). We excluded the Csp strain from this community, as it went extinct in 6 cases out of 8 in pairwise experiments (Fig. 2). These 8-strain experiments were performed six times independently. They showed that the 2 *Paraburkholderia* strains largely dominated these communities, since together they accounted for ≥ 95% of the final populations in four experiments (experiments 1, 2, 5 and 6) and 72% or 85% of the total populations in the other two experiments. It is interesting to note that Pphy alone represented ≥ 70% of the community in the four experiments showing the highest proportion of *Paraburkholderia* strains. Most *Cupriavidus* strains were not completely excluded, and their total and relative abundances fluctuated among experiments. Ctai5 and Ctai2 were rarely or never detected, respectively. Within each experiment, variability between replicates (each consisting of a pool of 3 plants) was moderate (Supplementary Fig. 6), suggesting that between-experiment variability may originate from inoculum preparation or small uncontrolled variations in plant culture conditions. Further analysis of the origin of this variability is beyond the scope of this study, but would represent a worthwhile follow-up of this work, as it was shown in other systems that small physiological differences between host or microbial individuals, or spatial structure of the environment, can lead to significant variations in the outcome of competitive infections [32–34]. To compare the results from the 8-strain communities to the pairwise inoculations, we first calculated a ‘dominance index’ in these 8-strain communities, similar to the one calculated for pairwise inoculations (Material and Methods). A positive correlation was observed between the two indexes (median rho = 0.93; 95% CI: 0.78 - 1.00; Spearman correlation test), showing that the hierarchy of strain dominance is conserved in the two types of experiments (Fig. 5D). High-ranking strains in the pairwise competitions (Pphy, Pcar, and to a lesser extent, Ctai1 and Ctai4) also robustly dominate in 8-strain communities. Similar results were obtained in a community comprising only the 6 *C. taiwanensis* strains. Indeed, Ctai1, Ctai4, Ctai6, and to a lesser extent Ctai3, were always recovered at relatively high frequency from these communities (Supplementary Fig. 7A and 8), but the dominant strain varied across replicate experiments (Ctai1, Ctai6 or Ctai4). Ctai5 was barely present while Ctai2 was always absent. Dominance indexes calculated in this community were also positively correlated with the dominance indexes calculated from pairwise inoculations (Supplementary Fig. 7B; median rho= 0.94; 95% CI: 0. 81-1.00; Spearman correlation test). Overall, these results show that symbiotic fitness is relatively robust and consistent across different levels of community complexity, and that higher-order interaction effects are either absent or weak in these communities.

**Figure 5.**
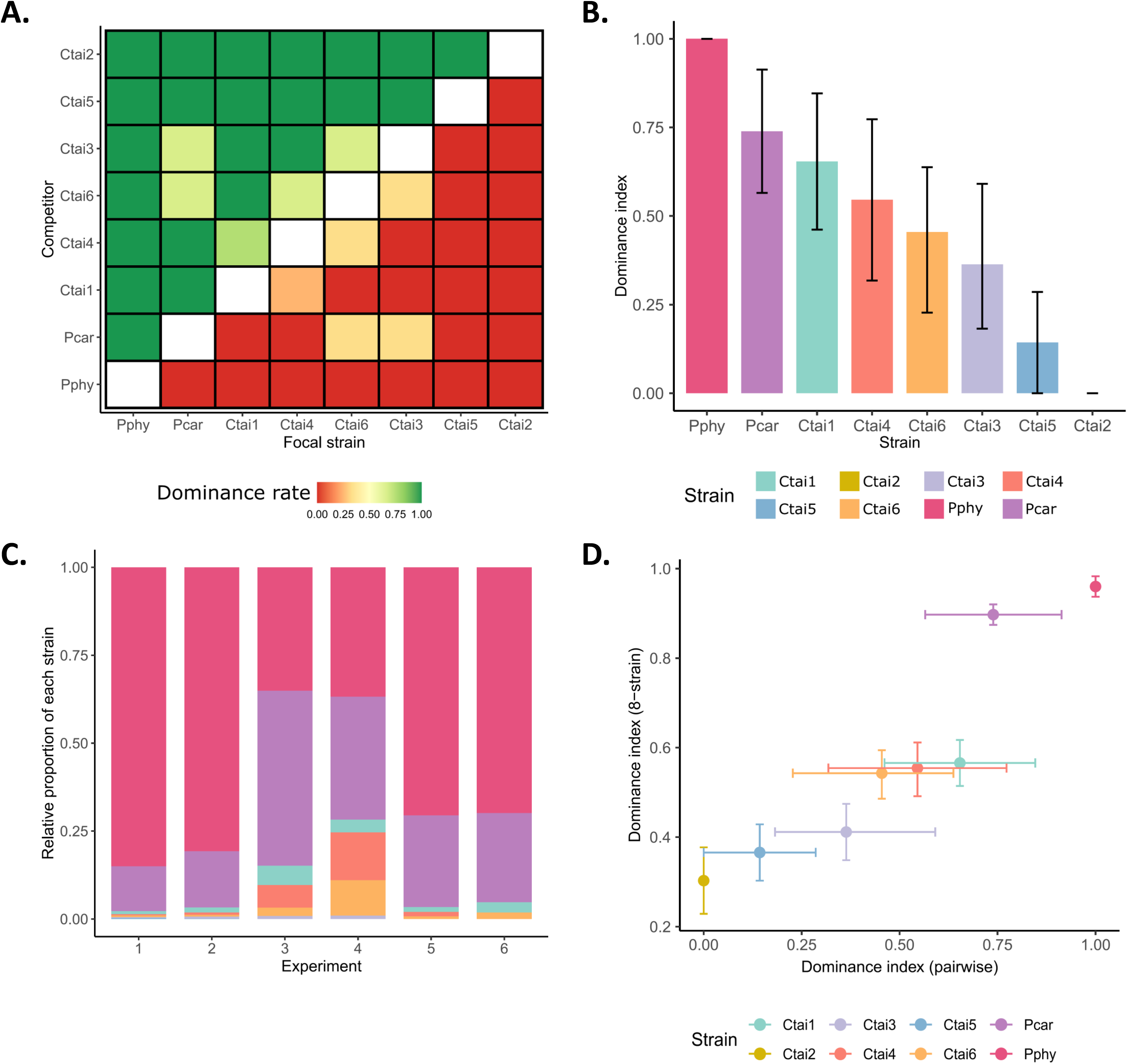
Transitivity in pairwise interactions is conserved in more complex communities. (A) Dominance rate of each focal strain in pairwise co-inoculation tests. For each pair of strains, the proportion of experiments (out of 3 or 4 independent experiments) in which the focal strain is more abundant than its competitor is shown as a color gradient. (B) Dominance index calculated as the average of dominance rates in all pairwise inoculation data. (C) Relative frequency of strain abundances in 8-strain communities. Data from 6 independent experiments are shown. (D) Dominance index from 8-strain communities plotted against dominance index from pairwise inoculations.

## Discussion

Within complex host-associated microbial communities, numerous factors influence the fitness of microbial members, including direct microbe-microbe interactions [24,25,35] and indirect host-mediated control mechanisms [2,14]. Here, our aim was to investigate how the presence of other rhizobia affects directly or indirectly the fitness of a focal strain during its interaction with a legume plant.

A systematic analysis of all 36 pairwise competitions between 9 natural symbionts of *M. pudica* showed that interactions affecting fitness are extremely common when several strains co-exist. In fact, measuring fitness in single inoculation assays only exceptionally predicted fitness for both strains in pairwise inoculations. These results support the view that multi-strain inoculations are needed for a pertinent evaluation of rhizobial fitness [28,36], as diverse rhizobial strains generally co-exist in soils [37]. Going further, analysing the different components of rhizobial fitness showed that nodulation competitiveness is the main source of interaction between rhizobia, being more frequent and generally stronger than post-infection effects. Again, this observation is consistent with previous results. Indeed, phenotypic diversity for nodulation competitiveness is common in rhizobia and underpins ecological success of strains [18,19]. Among *M. pudica* symbionts, *Paraburkholderia* strains, and *P. phymatum* STM815 (Pphy, in our study) in particular, are known to be highly competitive [38–42], even if they may be outcompeted by *Cupriavidus* strains under specific soil conditions (alkaline pH, presence of heavy metals) or on certain *M. pudica* genotypes [40,43–46]. Interestingly, the analysis of transcriptomic responses of *P. phymatum* STM815 and *C. taiwanensis* LMG19424 to root exudates of *M. pudica* showed that STM815 had a broader response than LMG19424 and included numerous genes known to be involved in plant-bacteria interactions [47]. These results suggest a better adaptation of STM815 to the rhizosphere of *M. pudica*, and are in line with the ancient evolutionary history of *P. phymatum* as a *Mimosa* symbiont [48]. Yet, the precise genetic bases of elevated nodulation competitiveness remain mostly unknown, as numerous bacterial traits and genetic determinants could be involved in this trait, including better survival and proliferation in the rhizosphere, higher motility, direct interference with competing strains or specific plant-bacterial recognition processes [18]. Moreover, whether differences in nodulation competitiveness arise from direct or indirect (*e.g.* mediated by the host plant) inter-bacterial competitions can be difficult to assess experimentally. While direct inter-bacterial interactions can be investigated in *in vitro* cultures [49] or in the rhizosphere (as performed in this study), these traits can only be correlated to nodulation competitiveness, unless genetic analysis is performed. Such broad scale genetic analyses of rhizobium competitiveness can include genome-wide association stuides [50], TnSeq [51] or experimental evolution coupled to whole-genome resequencing [52,53]. Alternatively, candidate gene approaches can prove informative. For example, the type-6 secretion system (T6SS) was shown to be a major determinant of nodulation competitiveness of *P. phymatum* STM815 against other *Paraburkholderia* strains, as well as an efficient antimicrobial weapon against these strains, strongly suggesting that direct antibacterial activity is responsible for high nodulation competitiveness [54]. However, in these experiments, the T6SS mutant remains a very good nodulation competitor, indicating that other properties of *P. phymatum* STM815 are involved in high competitiveness. These properties are likely important in our experiments, as we did not detect any consistent reduction in rhizospheric populations of rhizobia competing against *P. phymatum* STM815, although *P. phymatum* was extremely competitive in symbiosis against all strains tested.

A strong motivation for the detailed analysis of fitness in individual nodules of co-inoculated plants was the recent discovery of conditional sanctions in peas, which modifies bacterial loads within nodules depending on the relative levels of nitrogen provided to the plant by the two competing strains [22]. So far and to our knowledge, it is unknown if this mechanism can occur in plants other than pea, if it acts on natural strains’ diversity, nor if other types of post-infection indirect interactions can affect rhizobial fitness. Our experiments on *M. pudica* detected a few cases of such indirect interactions, but they were rare and their effect was generally minor compared to nodulation competitiveness. In one case (Ctai2-Csp), our results are consistent with a mechanism of conditional sanctions, since the fitness of the weakest mutualist, Csp, is decreased when co-inoculated with Ctai2. In another case (Ctai1-Ctai6), we ruled out the involvement of nitrogen fixation in the establishment of conditional post-infection control. Much work remains to be done to fully characterise this phenomenon (including the description of nodule cell infection, measuring bacterial proliferation and survival within nodules at different time points, or the quantification of resource supply to bacteroids), and identify the underlying molecular mechanisms. An interesting observation, however, is that the decrease in Ctai6’s fitness seems more drastic when considering CFU recovered from nodules than qPCR data (Fig. 4). This suggests that the viability of Ctai6 is severely altered during competition with Ctai1. Although *C. taiwanensis* cells are not terminally differentiated in *M. pudica* nodules [55], it is possible that some processes controlling bacterial viability and survival (*e.g* involving anti-microbial peptides or other defence responses, and that do not completely degrade DNA) are activated in these nodules [23]. In addition, it would be interesting to test if this phenomenon relies on a systemic signal. Many of these signals control different aspects of the rhizobium-legume symbiosis, including auto-regulation of nodulation, auto-regulation of infection or light-dependent control of nodulation [19,56,57]. Given that most of the molecular investigations concern single inoculations, we believe that many novel regulatory mechanisms remain to be discovered in cases where legume plants are co-inoculated with several rhizobial strains.

Finally, we observed a robust, transitive dominance hierarchy between our strains, both in pairwise inoculations and in more complex communities comprising 6 or 8 strains. This result indicates that, in our conditions, higher-order interactions (HOI) do not play a major role in determining community composition, and that fitness in pairwise competitions can help predict the composition of more complex communities. Understanding how microbe-microbe interactions shape complex microbial communities is a current major challenge for microbial ecology [58], with some studies detecting HOI effects [30,59–61] while others do not [62–65]. Interestingly, other examples of interactions among rhizobial communities were shown to be transitive interactions [66,67]. A future challenge for microbial ecology will be to understand why certain microbial communities are less likely to exhibit HOI than others, and to evaluate the role of eukaryotic hosts in these processes.

## Material and methods

### Plant growth

*Mimosa pudica* seeds were produced at LIPME in 2020 from a commercial seed lot sourced from B&T World Seed, Paguignan, France, originally of Australian origin. Seeds were sterilized as described in [68]. Seeds were first immersed in 95% H2SO4 during 15 minutes, rinsed in sterile distilled water 5 times, and then immersed in 2.4% sodium hypochlorite solution during 10 minutes and rinsed in sterile distilled water 5 times. Sterilized seeds were soaked in sterile water at 28°C under agitation for 24 hours and then transferred to soft agar and incubated in darkness at 28 °C for an additional 24 hours.

Germinated seedlings were transferred to glass culture tubes under nitrogen-free conditions. Each tube contained 20 mL of solid Fahraeus medium [69] and 40 mL of 1:4-diluted Jensen liquid medium [70].

Plants were cultivated in a growth chamber at 28 °C with a 16 h light / 8 h dark photoperiod. Tubes were randomly positioned within each experimental replicate to control for positional bias.

### Bacterial strains and growth conditions

All bacterial strains were cultivated at 28 °C on tryptone-yeast extract (TY) agar supplemented with 6 mM CaCl₂. The study included six strains of *Cupriavidus taiwanensis* (LMG19424 [Ctai1], STM6021 [Ctai2], STM8959 [Ctai3], STM8969 [Ctai4], STM8968 [Ctai5], STM8970 [Ctai6]); one *Cupriavidus* sp. strain amp6 (Csp); and two *Paraburkholderia* species (*P. phymatum* STM815 [Pphy], and *P. caribensis* Tj182 [Pcar]). Detailed strain information is provided in Table 1.

### Symbiotic phenotypes assays

For single inoculations, bacterial strains were grown on TY agar plates for 48 h and resuspended in sterile distilled water to an OD₆₀₀_nm_ of 0.02 (10^6^ CFU/mL). Each plantlet (3–4 days old) was inoculated with 100 μL of the bacterial suspension. To quantify *in planta* bacterial proliferation, nodules were harvested 28 days post-inoculation from three individual plants per treatment. Nodules were surface-sterilized (2.4% sodium hypochlorite, 15 min), rinsed with sterile distilled water, and homogenized by crushing with a sterile pestle in 1 mL of 5 mM MgSO₄. For shoot biomass measurements, shoot plant parts were dried at 65 °C for 48 h before weighing. To prepare samples for qPCR analyses, the nodule crush homogenate was vortexed and centrifuged at 100 × g for 5 min to remove root debris. From the supernatant, 100 µl were collected, boiled at 100°C for 15 min to lyse cells, cooled on ice for 5 min and stored at –20 °C.

### Pairwise inoculations

To assess relative symbiotic fitness, pairwise co-inoculations were performed using nine bacterial strains, resulting in 36 unique combinations. For each combination, *M. pudica* plantlets were inoculated with 100 μL of a mixed bacterial suspension containing equal proportions of both strains (1:1 ratio). To verify the initial inoculum proportion, serial dilutions of each bacterial suspension were plated on TY agar using an EasySpiral automatic plater (Interscience), and colonies were counted after 48 h at 28 °C.

Nodules were harvested 28 days post-inoculation from three individual plants, pooled, and crushed as described above. Bacterial strain abundance within the nodule homogenate was quantified by qPCR using strain-specific primers (Supplementary Table 1). Strain abundance was normalized by the proportion of each strain in the pooled nodules to its proportion in the initial inoculum.

### Nodulation competitiveness assays

Ten strain pairs showing coexistence were selected to assess nodulation competitiveness. The following combinations were tested: Ctai1–Ctai3, Ctai1–Ctai4, Ctai1–Ctai6, Ctai1–Pcar, Ctai2–Csp, Ctai2–Ctai5, Ctai3–Pcar, Ctai4–Ctai6, and Pphy–Pcar. For each combination, 93 individual nodules were isolated and crushed in flat-bottom 96-well plates. Controls were included in the remaining 3 wells: 2 wells contained one nodule from a single-inoculated plant with each strain of the tested pair and a negative control. Nodules were homogenized individually in 96-well plates in a total volume of 30 µl of 5 mM MgSO₄. From each homogenate, 10 µl was mixed with 90 µl of ultrapure water (Millipore), boiled for 15 min, cooled on ice for 5 min, and stored at-20°C before qPCR analysis. The identity of strain(s) present in each nodule was determined by qPCR using strain-specific primers, with each well tested independently.

### Individual nodule infectivity

The bacterial load within each nodule was assessed from individual nodules using qPCR quantification. In the case of Ctai1-Ctai6 and Ctai6-Ctai4 co-inoculations, viable bacteria per nodule was assessed by plating serial dilutions of nodule crushes on TY agar plates containing 6 mM CaCl2.

### Community inoculation assays

To evaluate strain interactions in a simplified community, we performed community-level inoculations by simultaneously inoculating *M. pudica* plantlets with a mixture of six (the six *Cupriavidus taiwanensis* strains) or eight strains (the six *Cupriavidus taiwanensis* strains, and the two *Parabrukholderia* strains Pphy and Pcar). The strain Csp was omitted from these experiments because it was absent at almost 100% of the pairwise inoculations. Plants were inoculated with equal proportions of each strain. Inoculation density and conditions were the same as in pairwise co-inoculations (100 μL total volume at 10^6^ CFU/mL). To verify the initial inoculum proportion, serial dilutions of each bacterial suspension were plated on TY agar using an EasySpiral automatic plater (Interscience), and colonies were counted after 48 h at 28 °C. Nodules were harvested 28 days post-inoculation from three individual plants per replicate, pooled, and crushed as described previously. Bacterial abundance within the nodule population was quantified by qPCR using strain-specific primers. As in pairwise assays, relative abundance was normalized to each strain’s initial proportion in the inoculum.

### Rhizosphere survival and colonisation assays

To assess rhizosphere colonization and survival, we inoculated plantlets with either single strains or 1:1 mixtures. Seven days post-inoculation, roots were harvested along with 10 mL of liquid Jensen medium and transferred into sterile 25 mL tubes. Samples were vortexed for 30 s to detach rhizosphere bacteria, after which roots were removed and the suspensions centrifuged at 8000 rpm for 5 min. Supernatant (9 mL) was discarded, and the bacterial pellet was resuspended in the remaining 1 mL. Samples were boiled at 100 °C for 15 min, cooled on ice for 5 min and stored at –20 °C for subsequent qPCR analysis.

### Design of primers, standard preparation and qPCR

Genomic DNA from each strain was extracted from liquid TY cultures using the Wizard® Genomic DNA Purification Kit (Promega). DNA concentration and purity were determined using both a NanoDrop™ spectrophotometer and a Qubit™ fluorometer. For each of the nine strains analyzed in this study, a pair of strain-specific oligonucleotides targeting the *gyrB* locus was designed according to standard quality criteria, including melting temperature (Tm), GC content, amplicon size, and sequence specificity (Supplementary Table 1).

To verify primer specificity and amplification efficiency, cross-amplification tests were performed for all primer pairs against all genomic DNA targets (50 ng/µl), and standard curves were generated from purified genomic DNA (Supplementary Figure 3A). Standard curves consisted of four 10-fold serial dilutions ranging from 2 ng/µl to 0.0002 ng/µl. Amplification specificity was confirmed by melt curve analysis at the end of each qPCR run. The slope, y-intercept, and coefficient of determination (R²) values of the standard curves are reported in Supplementary Figure 3B. Each 7 µl qPCR reaction contained 3.5 µl of Takyon™ No Rox SYBR® MasterMix (Eurogentec), primers at a final concentration of 10 µM each, and 2 µl of template DNA. Amplifications were carried out on a LightCycler® 480 system (Roche) under the following cycling conditions: initial denaturation at 95 °C for 3 min, followed by 40 cycles of 95 °C for 10 s, 58/62 °C for 15 s, and 72 °C for 15 s. When required, a melting curve program was included (95°C for 10 s, 55 °C for 15 s, followed by a gradual increase to 95 °C).In each qPCR run, a standard curve was included using 2 ng/µl of genomic DNA. To calculate DNA copy numbers, an average standard curve was generated for each strain using all standard curves obtained, along with the known genome size of each strain (Table 1).

For experimental assays, qPCR was performed on various sample types, including nodule homogenates (in single, pairwise, and multiple inoculation) and rhizosphere samples. Frozen samples (prepared as previously described in the respective sections) were thawed on ice. A 25-fold dilution was then prepared, and 2 µl of this diluted lysate was used as template for qPCR. When analysing qPCR results, we defined a threshold Ct value of 35, which corresponded on average (across all strains) to approximately 3.10⁴ DNA copies per plant. Because this quantity is below the expected bacterial content of a single nodule, samples with Ct ≥ 35 were considered below the detection threshold.

### Statistical analyses

All statistical analyses were performed in R version 4.2.2 [71], using the packages dplyr [72], tidyverse [73], lme4 [74], dunn.test [75] and ggplot2 [76].

### General modelling framework

Depending on the type and distribution of the response variables, we used either parametric tests, non-parametric tests or generalized linear (mixed) models (GLMMs). All GLMMs were defined with appropriate error distributions (binomial or gamma) and link functions (logit or log). To account for variability between experimental batches, we included a random intercept for the experimental batch in all models and where applicable, nested random effects to account for hierarchical experimental designs (e.g., plant identification nested within experimental batch). The significance of fixed effects was tested by comparing models with and without these fixed effects using likelihood ratio tests via the anova() function. Full model outputs are provided in the Supplementary Online Material.

### Symbiotic phenotype comparisons

Kruskal–Wallis rank-sum tests were used to assess variations in symbiotic phenotypes (shoot dry weight, nodule number, and bacterial abundance in nodules) across strains. When significant differences were detected, post hoc pairwise comparisons were conducted using Dunn’s test with Bonferroni correction to adjust for multiple testing.

### Pairwise combination screen

To evaluate inter-strain interactions in 36 pairwise combinations, we generated the distribution of predicted strain abundance under the null ‘no-interaction’ hypothesis by bootstrapping single inoculation data (n = 1000 iterations per strain, sampling with replacement). For each pair of strains, expected abundance of each strain in co-inoculation was calculated from randomly pooling three single-inoculation values (to take into account the fact that pairwise inoculation data came from 3 pooled plants while single inoculation data came from individual plants), and 95% confidence intervals (CIs) were computed. Observed co-inoculation values were compared to these CIs. Interaction outcomes were classified as dominance (only one strain detected) or coexistence (both strains detected). Within coexistence, changes in abundance, indicative of possible interactions between strains, were considered likely when all observed values fell outside the predicted CI from single inoculation data.

### Comparison of pairwise and community inoculations

A dominance index was calculated for each strain based on its performance in pairwise co-inoculations: we recorded whether a strain won or lost against its competitor. Winners were defined as the strain reaching a higher absolute abundance than its competitor and were assigned a score of 1; losers were assigned a score of 0. To account for experimental variability, we performed a bootstrap approach by resampling the co-inoculation replicates of each pairwise combination (n = 1000 iterations, sample with replacement). For each bootstrap iteration, the dominance index of a strain was defined as the mean score across all its pairwise competitions. A similar bootstrap procedure (n = 1000 iterations, sampling with replacement) was used to compute dominance indexes from the 6-strain and 8-strain community inoculations. Because these experiments were performed in batches, bootstraps were generated by sampling replicates within each batch to account for the experimental structure. In these community assays, observed values were obtained by the same qPCR-based quantification approach as in pairwise co-inoculations.

To assess whether dominance indexes measured in pairwise inoculations could explain the strains performance in more complex communities, we computed a Spearman rank correlation between the pairwise and community dominance indexes for each bootstrap iteration. This procedure generated a distribution of correlation coefficients, which we used to evaluate whether strain rank observed in pairwise inoculations were maintained in 6-strain or 8-strain communities.

### Phylogenetic analysis

DNA sequences from the *recA* genes were recovered from the following whole genome sequences available on the NCBI database: GCA_000069785.1 (*C. taiwanensis* LMG19424), GCA_900249815.1

(*C. taiwanensis* STM6021), GCA_900250045.1 (*C. taiwanensis* STM8959), GCA_900250015.1

(*C. taiwanensis* STM8969), GCA_900249975.1 (*C. taiwanensis* STM8970), GCA_900250125.1

(*C. taiwanensis* STM6160), GCA_000426345.1 (*Cupriavidus sp.* AMP6), GCA_000020045.1

(*P. phymatum* STM815) and GCA_003028645.1 (*P. caribensis* Tj182). Alignement was performed on translated sequences with MUSCLE [77]. The neighbour-joining phylogeny was computed with the BioNJ algorithm and 100 bootstraps with SeaView [78].

## Ackowledgements

This work is funded by the French National Research Agency (ANR-21-CE02-0019-01), the “Laboratoires d’Excellence (LABEX)” TULIP (ANR-10-LABX-41), and the “École Universitaire de Recherche (EUR)” TULIP-GS (ANR-18-EURE-0019). We thank Ludovic Mayran for help with preliminary experiments, and Marie Simonin and Robin Aguilée for insightful discussions on this project.

## Author contributions

MGA, BR, DC and PR designed the study. MGA, BR and PR performed the experimental work and analysed the data. JBF and DC contributed to data analysis. MGA and PR wrote the first draft of the manuscript. All authors contributed to the editing and finalizing of the manuscript.

## Data availability statement

All data and code generated during this work are available on the Data INRAE platform (https://doi.org/10.57745/TPHBT3).

Supplementary figure 1. Neighbour-joining phylogeny of RecA protein sequences from the 9 rhizobial strains used in this study.

Supplementary figure 2. Growth curves of the 9 rhizobial strains in TY medium. Points indicate the mean of four biological replicates, and error bars represent the standard deviation.

Supplementary figure 3. Validation of qPCR oligonucleotides used in this study. (A) Specificity of qPCR oligonucleotides tested against genomic DNA preparations of all rhizobial strains. Ct of each qPCR reaction is indicated in colored bins. (B) Efficiency of qPCR oligonucleotides tested against genomic DNA dilutions of their respective target strains. Equations and R2 values of linear fits of Ct values against logarithmic values of DNA concentrations are shown on each plot.

Supplementary figure 4. Comparison of symbiotic phenotypes in single and pairwise inoculations. (A) Number of nodules. (B) Shoot dry weight measured in *M. pudica* plants inoculated with the different bacterial strains and all their pairwise combinations. Measures were performed at 28 dpi and come from 4 independent experiments. n ≥ 8.

Supplementary figure 5. Rhizosphere colonisation. (A) Bacterial abundance in the rhizosphere during single inoculations. n ≥ 9. Kruskal-Wallis rank sum test with Dunn’s post-hoc correction. * indicates p-values < 0.05. (B) Correlation between relative bacterial abundance in the rhizosphere and nodulation occupancy during pairwise inoculations. Each point represents, for one focal strain of a given pair, the mean values of rhizosphere proportions and nodulation competitiveness measured in one experiment. n = 3.

Supplementary figure 6. Relative bacterial abundances in individual replicates from 6 independent experiments (E1-E6) inoculated with the 8-strain community. Each experiment comprised 4 or 5 replicate groups of 3 plants from which nodule bacterial populations were extracted and quantified by qPCR.

Supplementary figure 7. Symbiotic fitness in *C. taiwanensis* 6-strain community. (A) Mean relative frequencies of strain abundances in the 6-strain community. Data from 7 independent experiments are shown, each comprising 4 or 5 replicate groups of 3 plants. (B) Dominance index from 6-strain community plotted against dominance index from pairwise inoculations.

Supplementary figure 8. Relative frequencies of strain abundances in 6-strain communities. Data from 7 independent experiments (E1 to E7) are shown. Each experiment comprised 4 or 5 replicate groups of 3 plants from which nodule bacterial populations were extracted and quantified by qPCR.

