## Supplementary Material for "Transitive interactions among rhizobia determine their symbiotic fitness"

**SUPPLEMENTARY TABLE 1** | List of oligonucleotides.

| **Oligonucleotides** | **Sequences 5'-3'** | **%GC** | **tm** | **Description** |
|---|---|---|---|---|
| oCBM5942 | AATTTCTGACAGATCCGGTG | 45 | 58 | Reverse LGM19424 |
| oCBM5943 | GAAGTCGAACAGCAAACTGA | 45 | 58 | Forward LGM19424 |
| oCBM5944 | CGTGCGATGTTCGATCTCC | 58 | 60 | Reverse STM815 |
| oCBM5945 | CAGATAGCTCGTAGGCAAC | 53 | 58 | Forward STM815 |
| oCBM5946 | TCTGCTGGTCAAACGCTCT | 53 | 58 | Reverse STM6021 |
| oCBM5947 | TAAAGACTCCCATGTATGGC | 45 | 58 | Forward STM6021 |
| oCBM5948 | CTTTGGTGCGCTGGCTAC | 61 | 58 | Reverse STM6160 |
| oCBM5949 | CACCGATGGTCGTGATCG | 61 | 58 | Forward STM6160 |
| oCBM5950 | TGTCACGGTAGTTCAGCTG | 53 | 58 | Reverse STM8959 |
| oCBM5951 | TGAAGGCGTGGAGACTCG | 61 | 58 | Forward STM8959 |
| oCBM5952 | AGATTGTGGACTGGGAAGC | 53 | 58 | Reverse STM8968 |
| oCBM5953 | CCATGAGGAACATCAGGTC | 53 | 58 | Forward STM8968 |
| oCBM5954 | TGGTCGCATTATCCCAGAC | 53 | 58 | Reverse STM8969 |
| oCBM5955 | CGAATCTCTGTGAGTACCC | 53 | 58 | Forward STM8969 |
| oCBM5956 | CACTCCGTGCTCGTTGAC | 61 | 58 | Reverse STM8970 |
| oCBM5957 | GAAATCTATGCCAAGTCACTG | 43 | 60 | Forward STM8970 |
| oCBM5958 | GGAATGGATCGTGCTGGAG | 58 | 60 | Reverse TJ182 |
| oCBM5959 | ATACCTCGTAAGCCACGAC | 53 | 58 | Forward TJ182 |

### Supplementary Figure 1

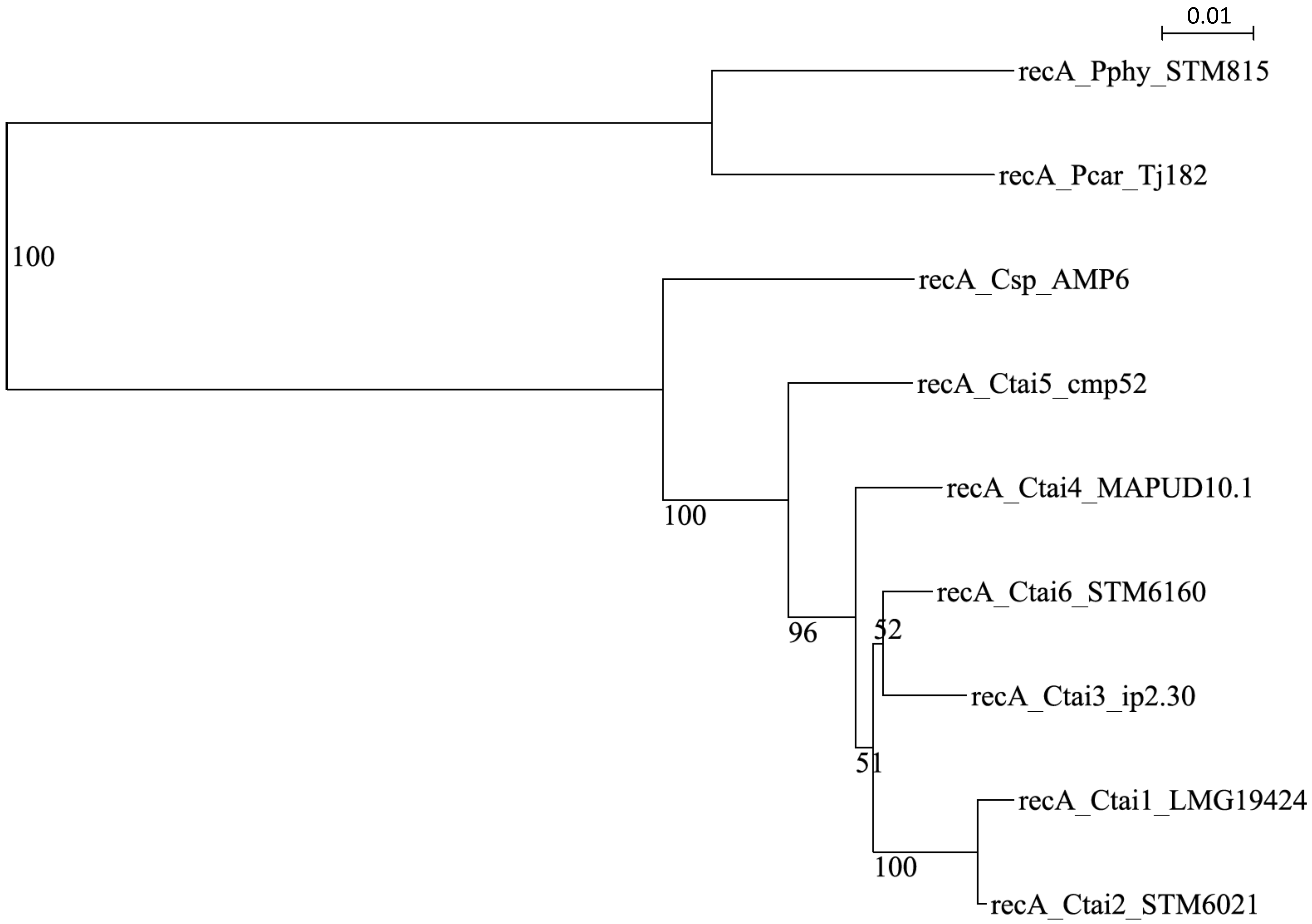

**Supplementary Figure 2**

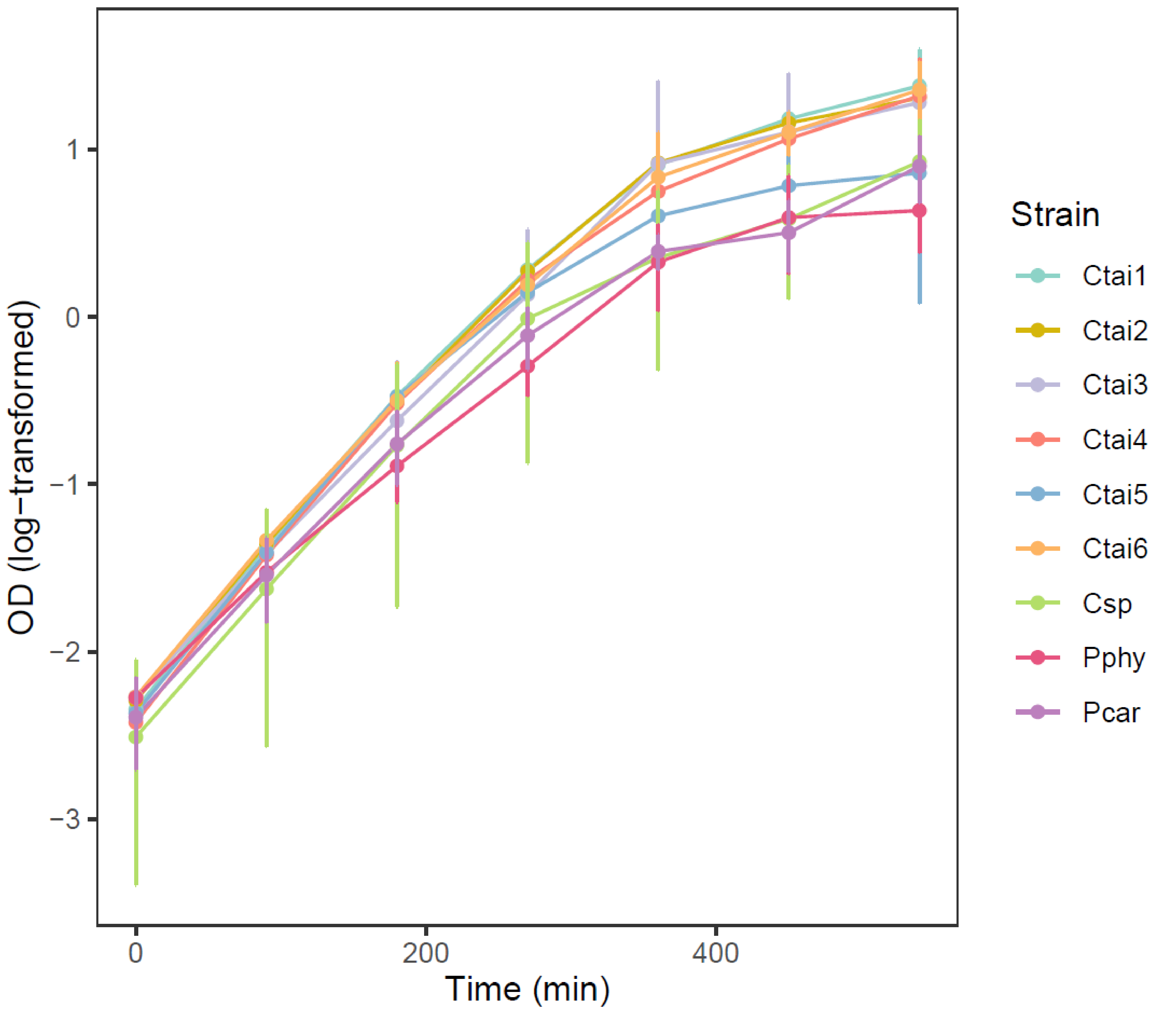

### Supplementary Figure 3

### A.

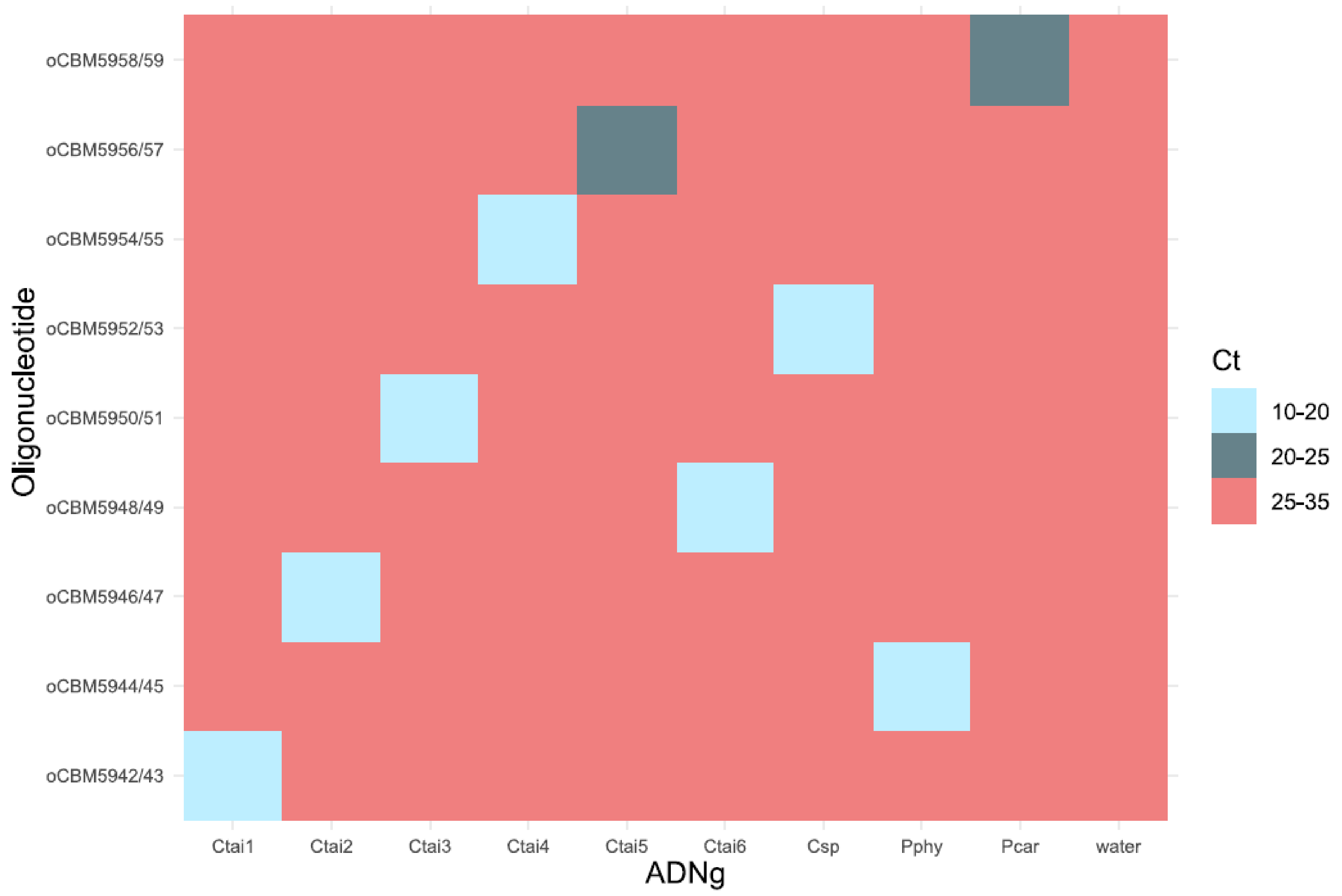


### B.

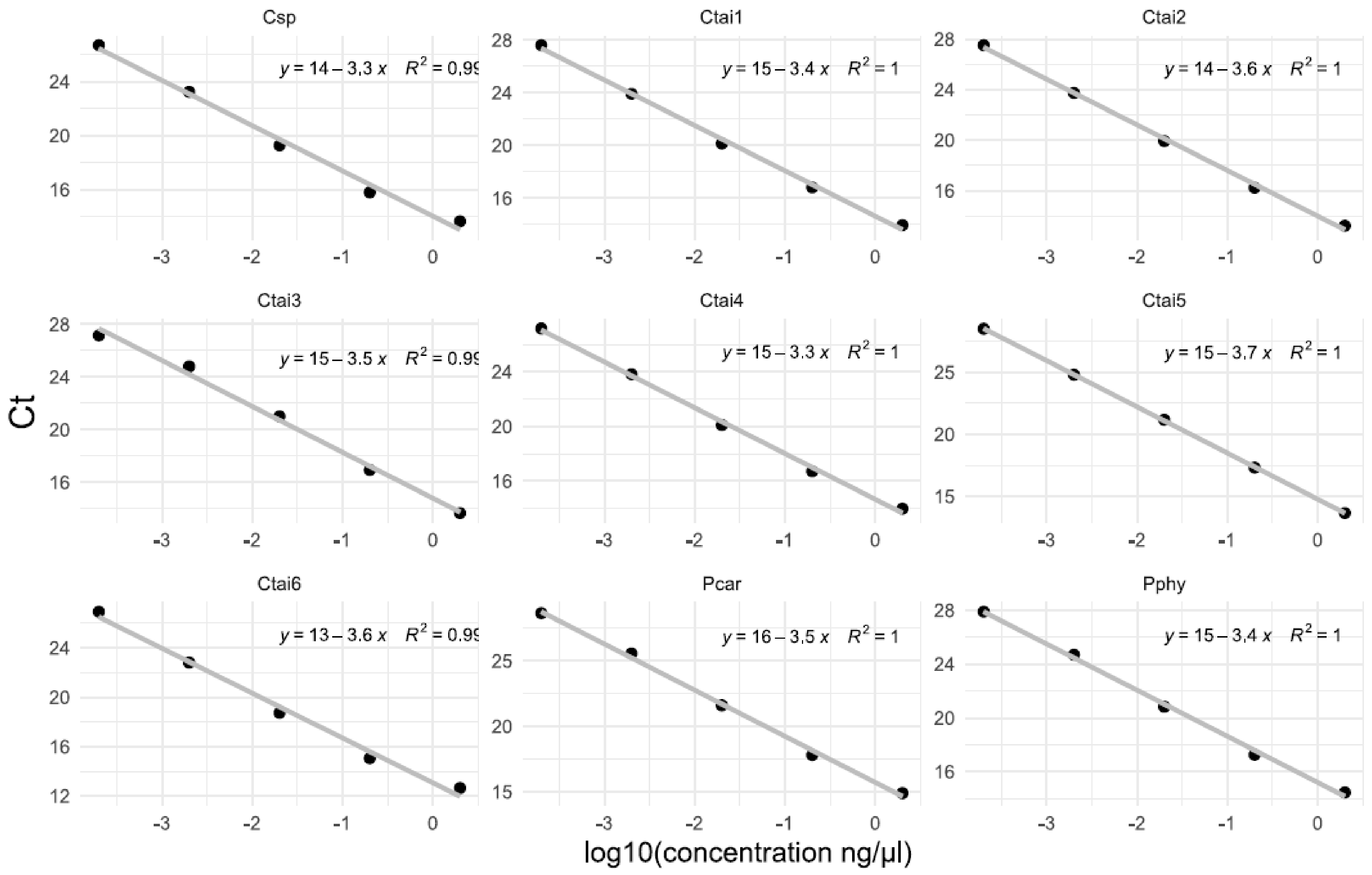

**Supplementary Figure 4**

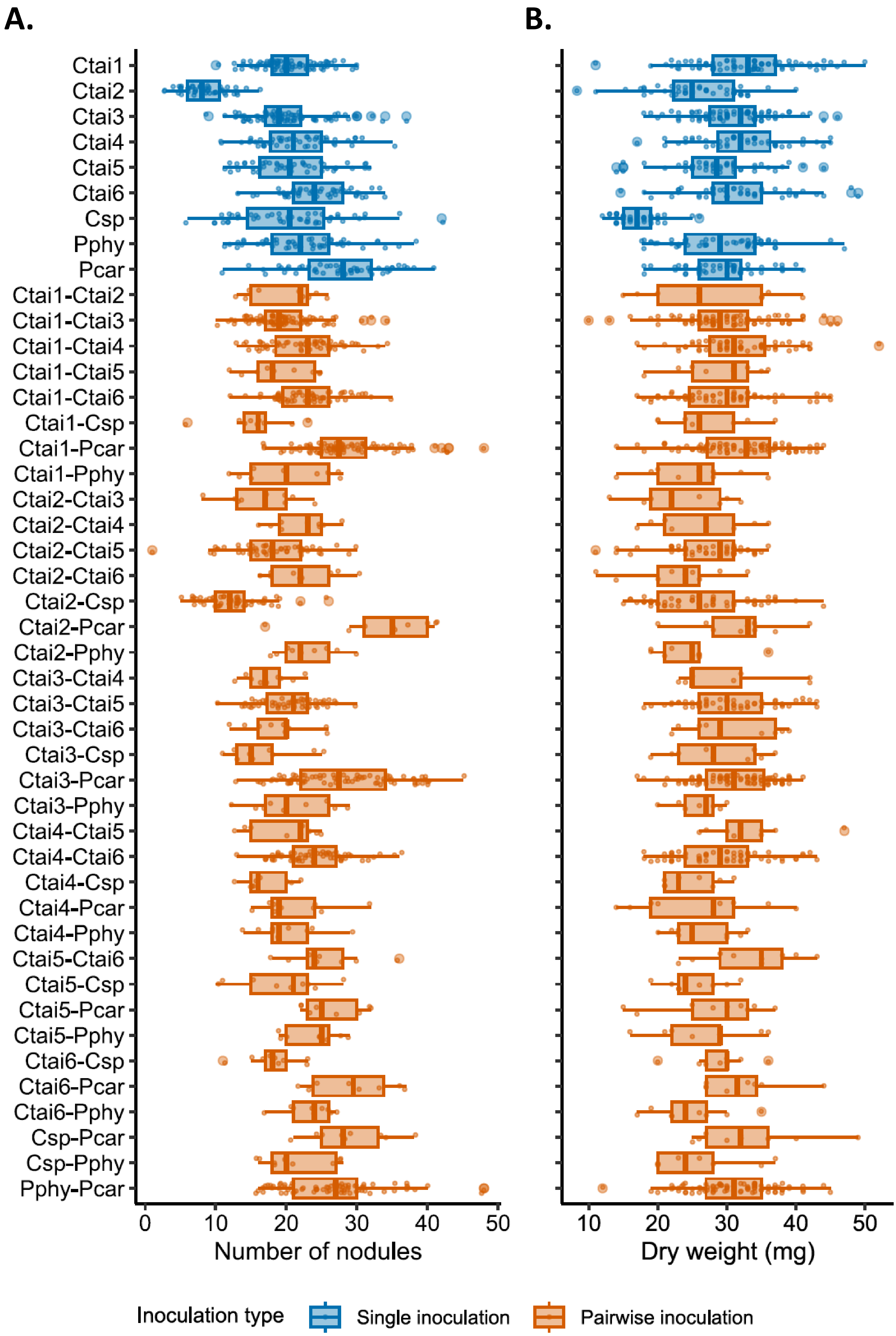

### Supplementary Figure 5

A.

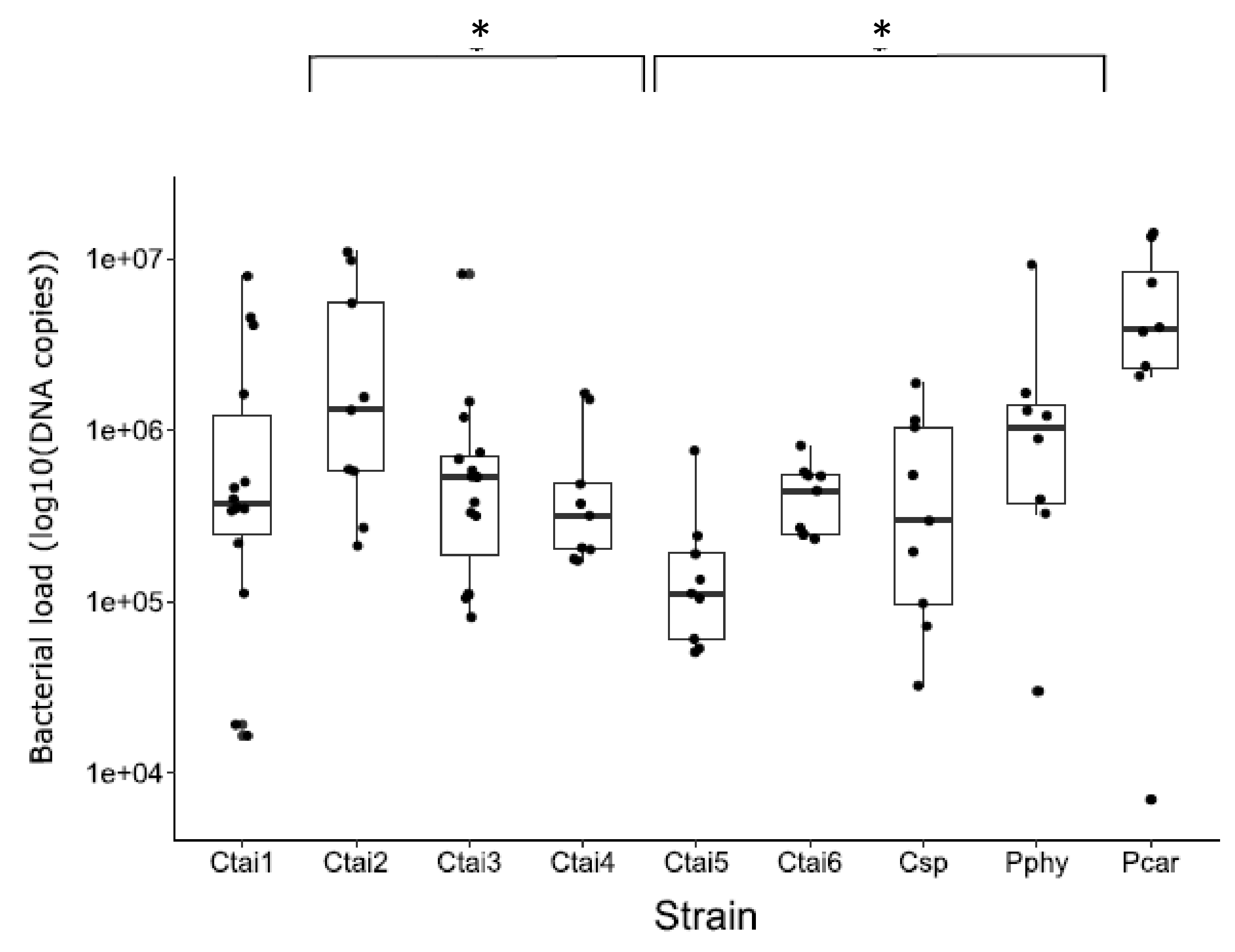


B.

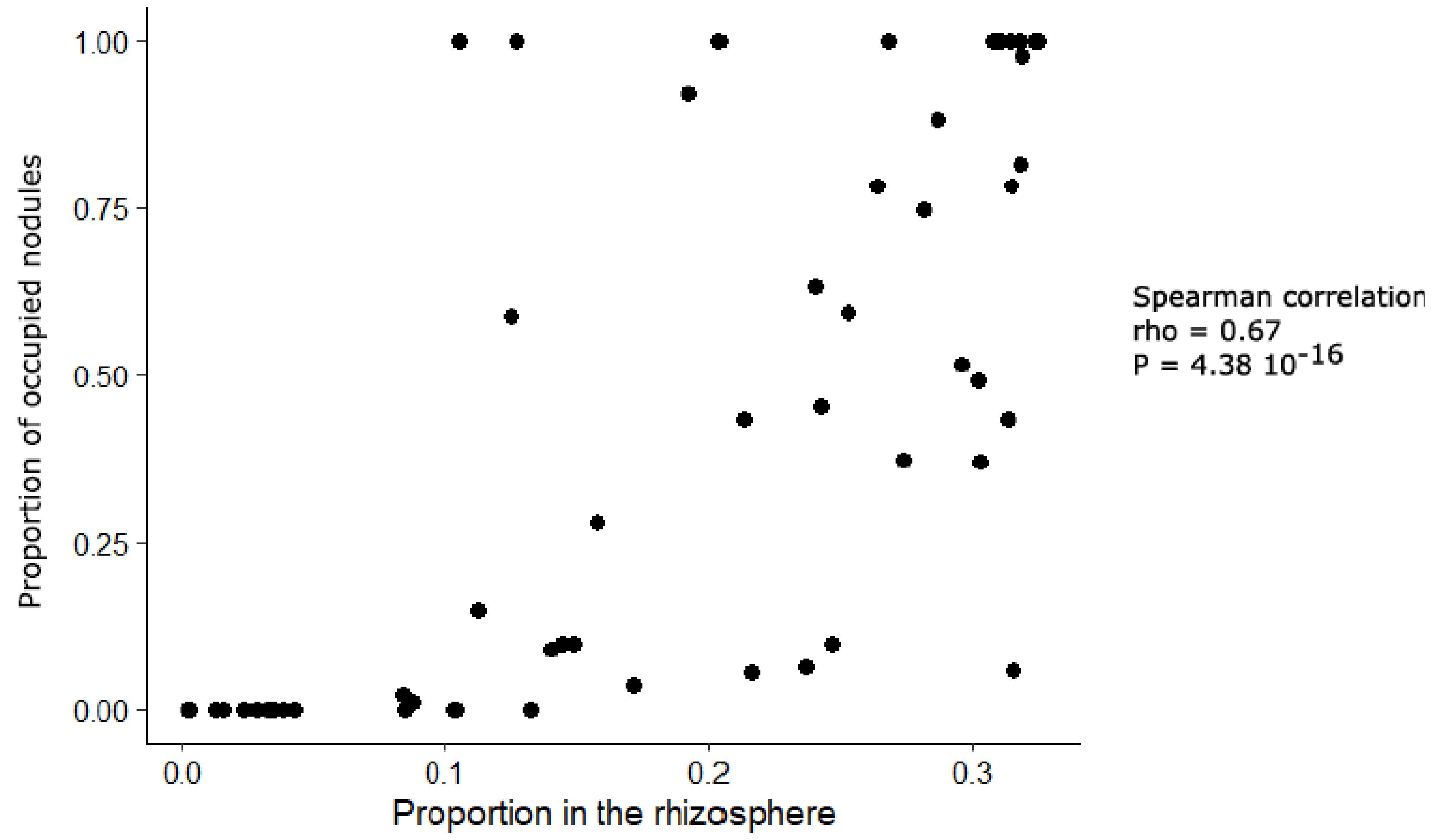

### Supplementary Figure 6

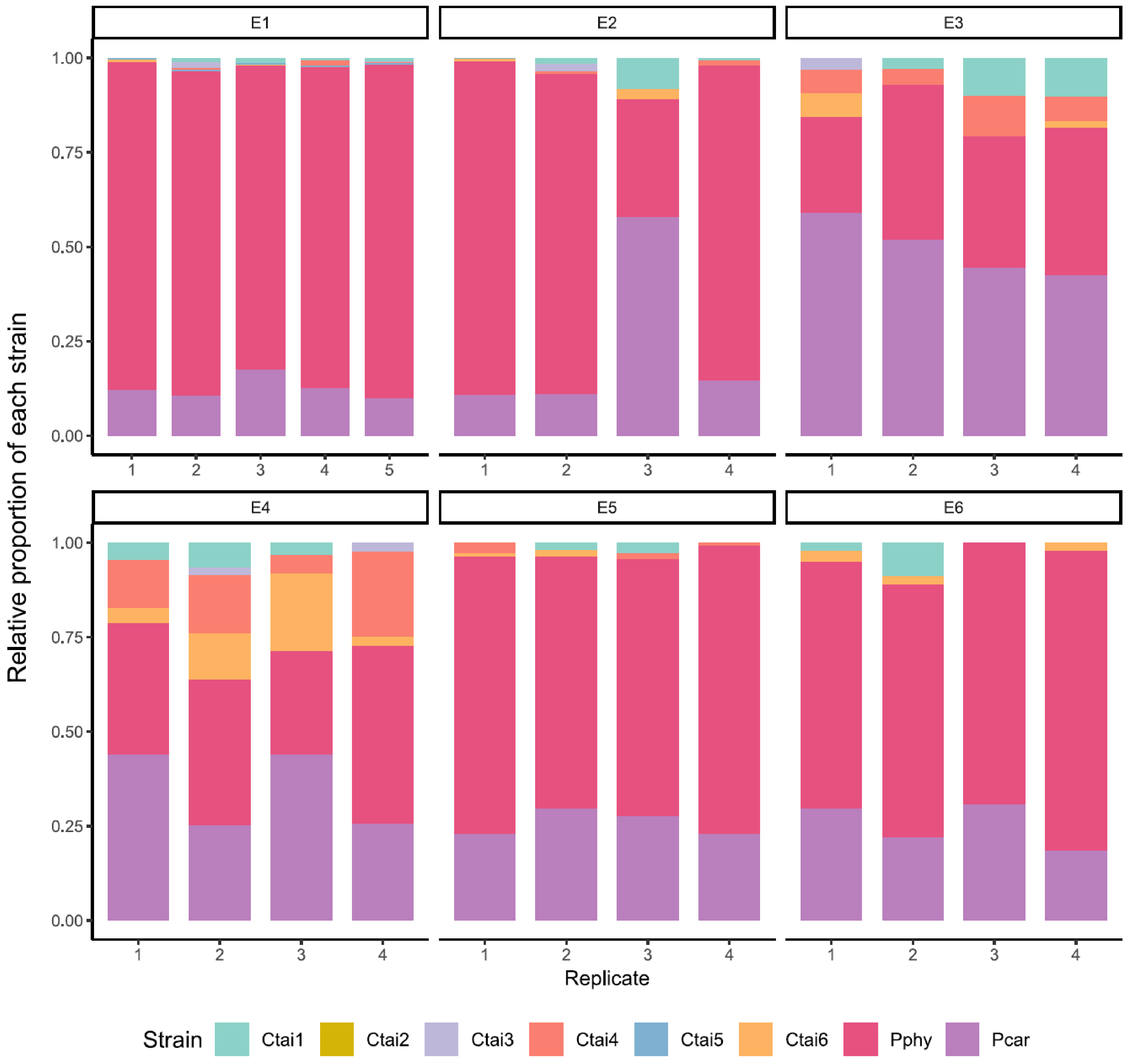

### Supplementary Figure 7

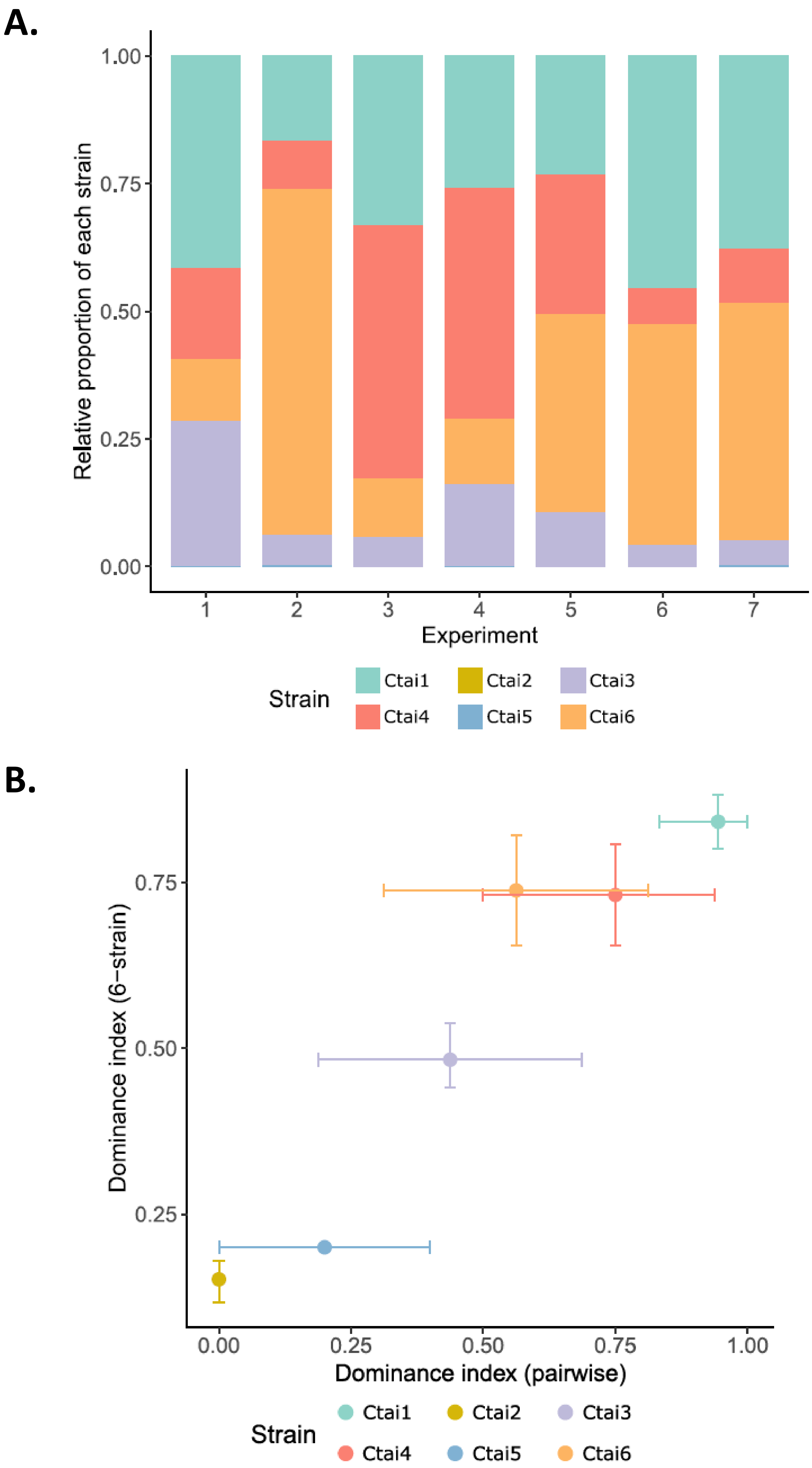

### Supplementary Figure 8

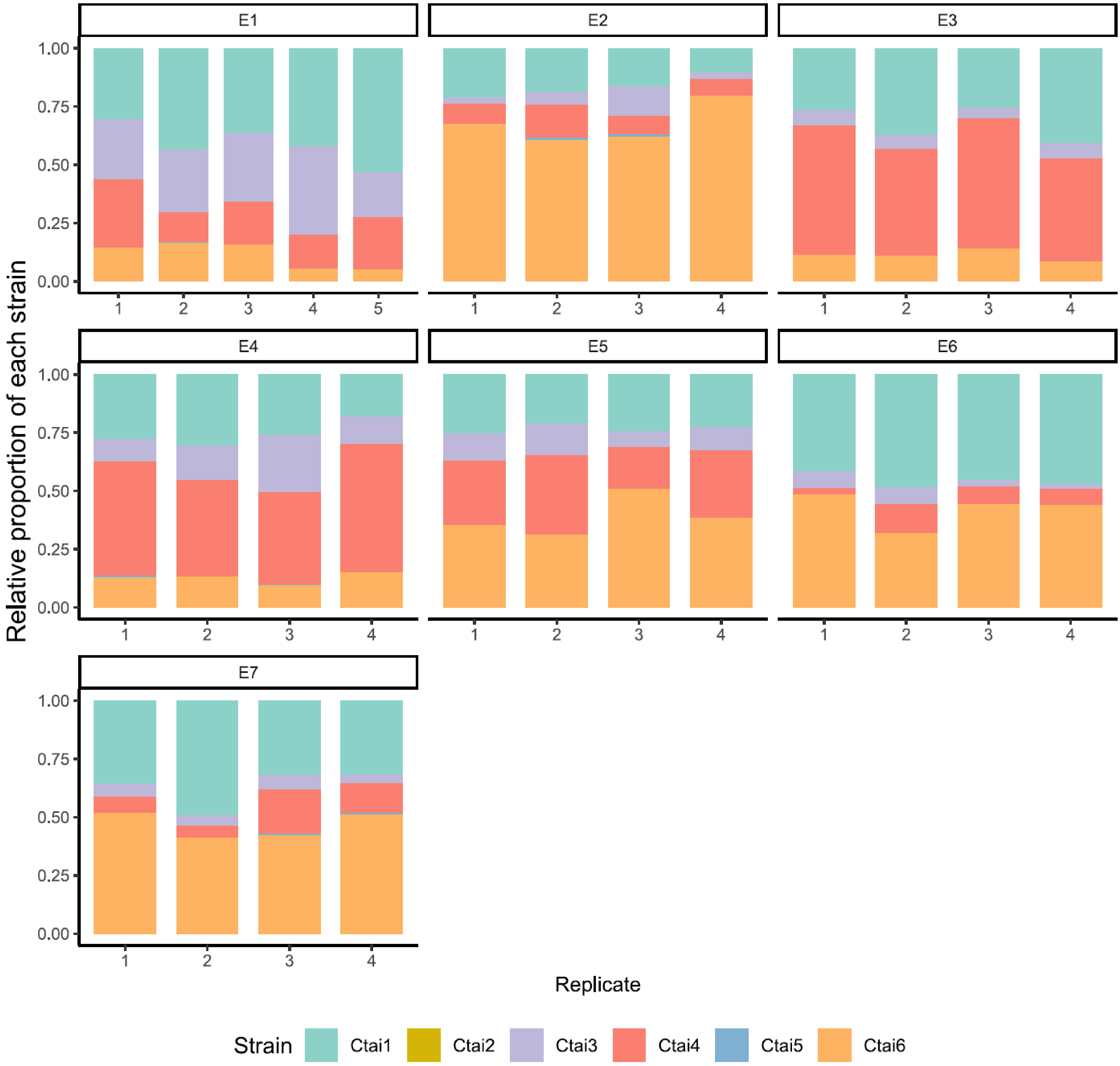

Supplementary online material (SOM) for the manuscript:

**Transitive interactions among rhizobia determine bacterial fitness during their interaction with the legume plant *Mimosa pudica***

Authors:

Margarita Granada Agudelo, Bryan Ruiz, Jean-Baptiste Ferdy, Delphine Capela, Philippe Remigi

This document presents the detailed outputs for statistical analyses performed in the manuscript. It contains 2 sections.

SOM section 1: Model estimates of the statistical model fitted to the nodule occupancy data presented in figure 3A

A binomial GLMM was fitted to model the proportion of nodules occupied by each strain in pairwise co-inoculations, using the sample (competing strains) as a fixed effect and a random intercept for experiment batch (glmer(), lme4).

Model: `glmer(formula = strain1_success ~ Sample_abb + (1 | Expt), family = binomial(link = "logit"))`

`Signif. codes:  0 '***' 0.001 '**' 0.01 '*' 0.05 '.' 0.1 ' ' 1`

Fixed effect

| | *Estimate* | *Std. error* | *z value* | *Pr(>\|z\|)* | |
|---|---|---|---|---|---|
| *(Intercept)* | -0.191 | 0.183 | -1.045 | 0.296 | |
| *Ctai1-Ctai3* | 0.592 | 0.244 | 2.422 | 0.015 | * |
| *Ctai1-Ctai4* | 0.755 | 0.165 | 4.571 | <0.001 | *** |
| *Ctai1-Pcar* | -1.182 | 0.231 | -5.122 | <0.001 | *** |
| *Ctai2-Csp* | 1.179 | 0.303 | 3.888 | <0.001 | *** |
| *Ctai5-Csp* | -2.792 | 0.386 | -7.231 | <0.001 | *** |
| *Ctai3-Ctai5* | 2.894 | 0.361 | 8.016 | <0.001 | *** |
| *Ctai3-Pcar* | -3.161 | 0.383 | -8.246 | <0.001 | *** |
| *Ctai4-Ctai6* | 0.261 | 0.176 | 1.480 | 0.139 | |
| *Pphy-Pcar* | 0.056 | 0.215 | 0.260 | 0.795 | |

Random effect

| *Group* | *Effect* | *Variance* | *Std. Dev* |
|---|---|---|---|
| *Expt* | Intercept | 0.101 | 0.318 |

SOM section 2: Model estimates of the statistical model fitted to the abundance in single nodule presented in figure 3B.

For each strain a GLM with gamma distribution and log link was used to compare strain infectivity at the single-nodule level in single vs. co-inoculations for each strain, using the sample (either single inoculation or competing strains) as a fixed effect and a random intercept for experiment batch (glmer(), lme4).

Model: `glmer(formula = Cp_ml ~ Sample_abb + (1 | Expt), family = Gamma(link = "log"))`

`Signif. codes:  0 '***' 0.001 '**' 0.01 '*' 0.05 '.' 0.1 ' ' 1`

Ctai1:

Fixed effect

| | *Estimate* | *Std. error* | *t value* | *Pr(>\|z\|)* | |
|---|---|---|---|---|---|
| ***(Intercept)*** | 16.21260 | 0.20727 | 78.221 | < 0.001 | *** |
| ***Ctai1-Ctai3*** | -0.28737 | 0.24063 | -1.194 | 0.23238 | |
| ***Ctai1-Ctai4*** | -0.05733 | 0.24321 | - 0.236 | 0.81366 | |
| ***Ctai1-Ctai6*** | 0.07131 | 0.24451 | 0.292 | 0.77056 | |
| ***Ctai1-Pcar*** | 0.73982 | 0.26010 | 2.844 | 0.00445 | ** |

Random effect

| *Group* | *Effect* | *Variance* | *Std. Dev* | *Levels* |
|---|---|---|---|---|
| *Expt* | Intercept | 0.0165 | 0.129 | 7 |
| *Residual* | | 0.2689 | 0.519 | 500 obs |

Ctai2:

| | *Estimate* | *Std. error* | *t value* | *Pr(>\|z\|)* | |
|---|---|---|---|---|---|
| ***(Intercept)*** | 16.75300 | 0.09513 | 176.107 | <0.001 | *** |
| ***Ctai2-Ctai5*** | 0.18294 | 0.15996 | 1.144 | 0.253 | |
| ***Ctai2-Csp*** | -0.03961 | 0.10968 | -0.361 | 0.718 | |

Random effect

| *Group* | *Effect* | *Variance* | *Std. Dev* | *Levels* |
|---|---|---|---|---|
| *Expt* | Intercept | 0.001544 | 0.03929 | 4 |
| *Residual* | | 0.169593 | 0.41182 | 201 obs |

Ctai3:

Fixed effect

| | *Estimate* | *Std. error* | *t value* | *Pr(>\|z\|)* | |
|---|---|---|---|---|---|
| ***(Intercept)*** | 15.72165 | 0.2733 | 57.516 | <0.001 | *** |
| ***Ctai1-Ctai3*** | 0.08834 | 0.31455 | 0.281 | 0.779 | |
| ***Ctai3-Ctai5*** | -0.01115 | 0.32049 | -0.035 | 0.972 | |
| ***Ctai3-Pcar*** | 0.33157 | 0.34063 | 0.973 | 0.330 | |

Random effect

| *Group* | *Effect* | *Variance* | *Std. Dev* | *Levels* |
|---|---|---|---|---|
| ***Expt*** | (Intercept) | 0.02194 | 0.1481 | 7 |
| ***Residual*** | | 0.20260 | 0.4501 | 398 obs |

Ctai4:

Fixed effect:

| | *Estimate* | *Std. error* | *t value* | *Pr(>\|z\|)* | |
|---|---|---|---|---|---|
| ***(Intercept)*** | 15.8418 | 0.2524 | 62.768 | <0.001 | *** |
| ***Ctai1-Ctai4*** | -0.1110 | 0.2960 | -0.375 | 0.708 | |
| ***Ctai4-Ctai6*** | 0.1387 | 0.2954 | 0.469 | 0.639 | |

Random effect

| *Group* | *Effect* | *Variance* | *Std. Dev* | *Levels* |
|---|---|---|---|---|
| ***Expt*** | (Intercept) | 0.01651 | 0.1285 | 4 |
| ***Residual*** | | 0.15435 | 0.3929 | 265 obs |

Ctai5:

Fixed effect

| | *Estimate* | *Std. error* | *t value* | *Pr(>\|z\|)* | |
|---|---|---|---|---|---|
| ***(Intercept)*** | 16.0447 | 0.1055 | 152.125 | <0.001 | *** |
| ***Ctai3-Ctai5*** | 0.1235 | 0.1217 | 1.015 | 0.310 | |
| ***Ctai5-Csp*** | 0.1511 | 0.1499 | 1.008 | 0.313 | |

Random effect

| *Group* | *Effect* | *Variance* | *Std. Dev* | *Levels* |
|---|---|---|---|---|
| ***Expt*** | (Intercept) | 0.003218 | 0.05673 | 4 |
| ***Residual*** | | 0.139341 | 0.37328 | 304 obs |

Ctai6:

| | Estimate | Std. error | t value | Pr(>\|z\|) | |
|---|---|---|---|---|---|
| ***(Intercept)*** | 15.7456 | 0.1617 | 97.380 | <0.001 | *** |
| ***Ctai1-Ctai6*** | -0.4122 | 0.1899 | -2.171 | 0.030 | * |
| ***Ctai4-Ctai6*** | -0.1355 | 0.1900 | -0.713 | 0.476 | |

Random effect

| Group | Effect | Variance | Std. Dev | Levels |
|---|---|---|---|---|
| *Expt* | (Intercept) | 0.01086 | 0.1042 | 4 |
| *Residual* | | 0.28803 | 0.5367 | 307 obs |

Pphy:

| | Estimate | Std. error | t value | Pr(>\|z\|) | |
|---|---|---|---|---|---|
| ***(Intercept)*** | 16.8678 | 0.6514 | 25.897 | <0.001 | *** |
| ***Pphy-Pcar*** | -0.7607 | 0.7494 | -1.015 | 0.31 | |

Random effect

| Group | Effect | Variance | Std. Dev | Levels |
|---|---|---|---|---|
| *Expt* | (Intercept) | 0.1293 | 0.3596 | 4 |
| *Residual* | | 0.2818 | 0.5308 | 169 obs |

Pcar:

| | Estimate | Std. error | t value | Pr(>\|z\|) | |
|---|---|---|---|---|---|
| ***(Intercept)*** | 16.2950 | 0.1758 | 92.715 | <0.001 | *** |
| ***Ctai1-Pcar*** | -0.1408 | 0.2028 | -0.694 | 0.487617 | |
| ***Ctai3-Pcar*** | -0.1885 | 0.2021 | -0.932 | 0.351187 | |
| ***Pphy-Pcar*** | -0.7578 | 0.2063 | -3.674 | <0.001 | *** |

Random effect

| Group | Effect | Variance | Std. Dev | Levels |
|---|---|---|---|---|
| *Expt* | (Intercept) | 0.01833 | 0.1354 | 4 |
| *Residual* | | 0.67097 | 0.8191 | 684 obs |
